# Enigmatic missense mutations can cause disease via creation of *de novo* nuclear export signals

**DOI:** 10.1101/2024.04.24.590854

**Authors:** By Michael McConville, Toby Thomas, Ryan Beckner, Catherine Valadez, YuhMin Chook, Stephen Chung, Glen Liszczak

**Affiliations:** Department of Biochemistry, University of Texas Southwestern Medical Center, Dallas, TX, USA; Department of Internal Medicine, University of Texas Southwestern Medical Center, TX, USA; Department of Pharmacology, University of Texas Southwestern Medical Center, Dallas, TX, USA

## Abstract

Disease-causing missense mutations that occur within structurally and functionally unannotated protein regions can guide researchers to new mechanisms of protein regulation and dysfunction. Here, we report that the thrombocytopenia-, myelodysplastic syndromes-, and leukemia-associated P214L mutation in the transcriptional regulator ETV6 creates an XPO1-dependent nuclear export signal to cause protein mislocalization. Strategies to disrupt XPO1 activity fully restore ETV6 P214L protein nuclear localization and transcription regulation activity. Mechanistic insight inspired the design of a ‘humanized’ ETV6 mice, which we employ to demonstrate that the germline P214L mutation is sufficient to elicit severe defects in thrombopoiesis and hematopoietic stem cell maintenance. Beyond ETV6, we employed computational methods to uncover rare disease-associated missense mutations in unrelated proteins that create a nuclear export signal to disrupt protein function. Thus, missense mutations that operate through this mechanism should be predictable and may suggest rational therapeutic strategies for associated diseases.

## Introduction

Disease-associated missense mutations commonly occur in protein regions that lack structural or functional epitopes^1,2^. Given their location within unannotated protein segments, the mechanisms whereby these mutations impact protein function are often difficult to predict and poorly understood. Efforts to comprehensively define how enigmatic missense mutations alter protein function can elucidate new modes of protein dysregulation, guide development of associated disease models, and lay the groundwork for rational therapeutic design^3^.

The transcriptional repressor protein ETV6 plays a critical role in hematopoiesis, including hematopoietic stem cell (HSC) maintenance, thrombopoiesis, and lymphopoiesis^4,5^. Many unique *ETV6* genetic alterations have been characterized as critical contributors to myeloid and lymphoid leukemogenesis^6–13^. Given their pervasive nature and high penetrance, intense efforts to elucidate how ETV6 mutations lead to protein dysfunction and disease phenotypes are ongoing. To date, more than 30 cancer-associated chromosomal translocation events involving *ETV6* have been reported^13^. While these translocations can create chimeric transcription factors or tyrosine kinases that are critical for transformation, it is often loss of ETV6 that is a key event in disease onset.

Consistently, loss-of-function somatic *ETV6* missense mutations are recurrently observed in MDS and leukemia sequencing analyses^8,9,14,15^. More recently, familial inherited autosomal dominant *ETV6* missense mutations have been reported, and carriers segregate with variable thrombocytopenia and an elevated risk for MDS and leukemia^16–18^. These disease associations are in line with phenotypes observed in conditional *ETV6* knockout mice, which demonstrate a role for ETV6 in adult HSC survival and megakaryocyte maturation^4,5^.

There are two ‘hotspots’ within ETV6 where somatic and germline missense mutations commonly occur^19,20^. The first hotspot is the ETS family DNA-binding (ETS) domain, wherein a cluster of mutations occur that disrupt the DNA:protein interaction^19^. The second hotspot is the P214 site, which lies within a long, unstructured linker that connects the N-terminal Pointed (PNT) domain to the ETS domain^16,18^. This site does not fall within a functionally annotated region of ETV6 and the mechanism whereby a P214 mutation abrogates ETV6 activity is unclear. In the case of *ETV6^P214L^* germline mutations, all carriers are heterozygous for the allele and thrombocytopenic, and approximately 30% will develop leukemia in their lifetime^20^. Similar to many ETS domain mutations, including R369Q, R399C, R418G, the P214L mutation causes ETV6 mislocalization to the cytoplasm^16,18^. It has therefore been hypothesized that the P214L mutation abrogates DNA binding via an allosteric mechanism and that loss of DNA binding activity causes ETV6 cytoplasmic accumulation^18^. However, we found that the P214L mutation has no effect on the ETV6:DNA interaction in biochemical assays (Figure S1). Consistently, AlphaFold prediction of the full-length ETV6 structure does not suggest P214 interaction with the ETS or PNT domains (Figure 1A). Furthermore, mice carrying the homologous *Etv6^P216L/WT^* or *Etv6^P216L/P216^* allele do not exhibit overt hematopoietic defects^21^. These confounding observations have raised questions about the P214L mutation and its role in ETV6 dysfunction and hematologic disease.

**Figure 1.**
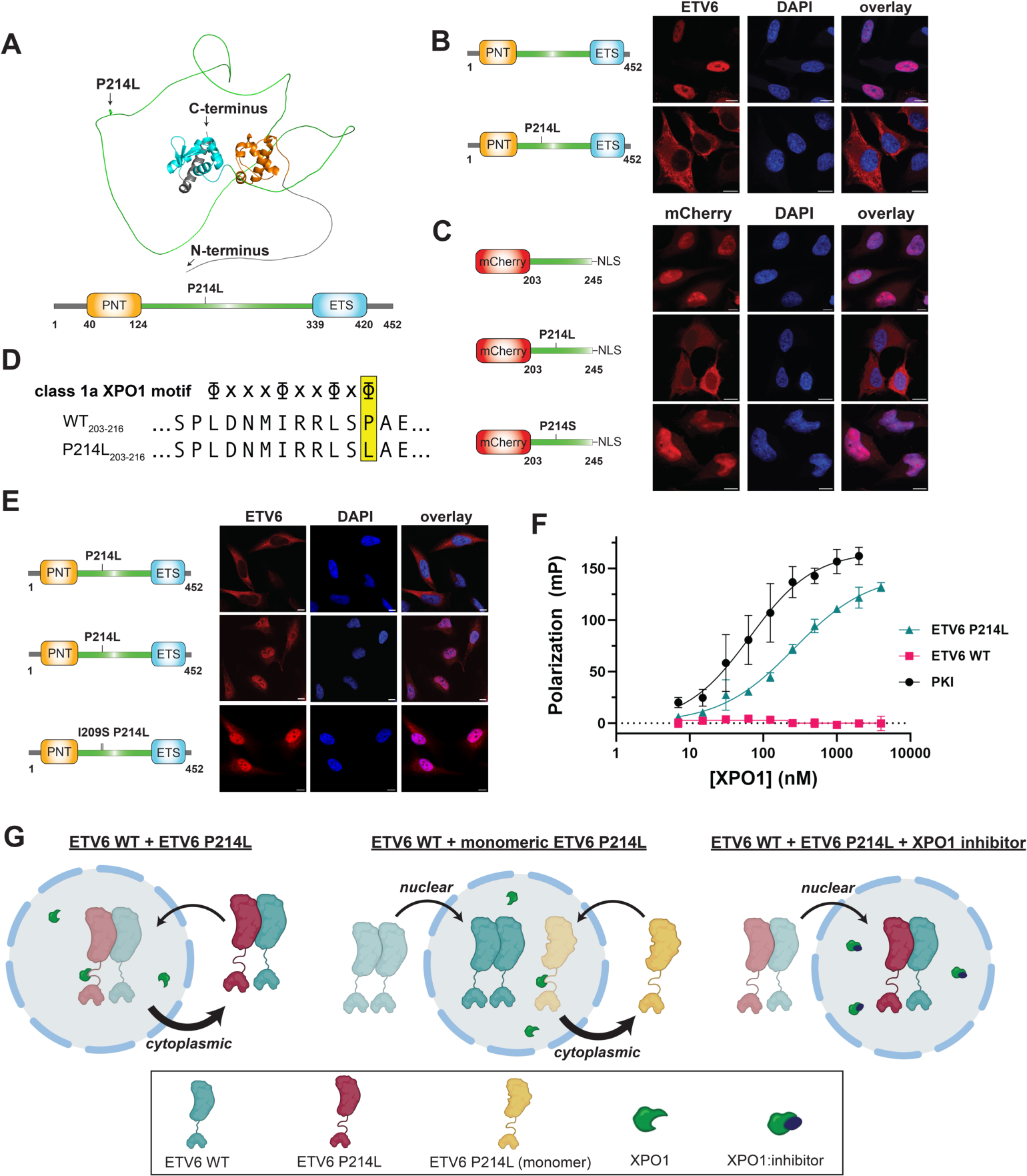
The ETV6 P214L mutation creates an NES to cause protein mislocalization. (A) The domain layout and AlphaFold-predicted structure of full-length ETV6. (B) ETV6 immunofluorescence microscopy z-stack images of HeLa cells bearing exogenous wild-type and mutant ETV6 constructs. Scale bar = 10 μm. (C) Fluorescence confocal microscopy z-stack images of HeLa cells bearing mCherry-fused ETV6-NLS fragments (amino acids 203-245). Scale bar = 10 μm. (D) Alignment of wild-type ETV6 and P214L mutant fragments overlaid with the class 1a XPO1 interaction motif. Φ = Ile, Leu, Met, Phe, or Val and X = any amino acid. (E) ETV6 immunofluorescence microscopy z-stack images of HeLa cells bearing exogenous ETV6 P214L protein in the absence and presence of Leptomycin B (5 nM) or the ETV6 I209S/P214L double mutant construct. Scale bar = 10 μm. (F) A fluorescence polarization interaction assay to assess binding of synthetic, fluorescently-labeled wild-type and mutant ETV6 fragments (amino acids 203-245) to a reconstituted XPO1:RanGTP nuclear export receptor complex. A fluorescently labeled variant of the strong XPO1 ligand peptide PKI is included. Error bars represent S.D. from n = 3 experimental replicates. (G) A schematic summarizing the mechanistic basis for the dominant negative effect of the ETV6 P214L mutation. Note: ‘monomeric ETV6’ = ETV6 construct bearing A93D and V112E mutations, which block PNT domain oligomerization. For corresponding immunofluorescence imaging experiments, see Figure S5.

Here, we set out to understand how the enigmatic P214L mutation disrupts ETV6 localization and function. We have discovered that the P214L mutation creates a potent nuclear export signal (NES) and that ETV6 mislocalization is dependent upon the activity of the nuclear export receptor, XPO1. This mechanism explains the dominant negative *ETV6^P214L/WT^* phenotype, as wild-type and mutant ETV6 protein copies oligomerize via the PNT domain resulting in nuclear export of both constructs. We also find that strategies to disrupt the mutant ETV6:XPO1 interaction restore ETV6 nuclear localization, rescue ETV6 transcription regulation activity, and restore Mendelian inheritance patterns in mice. We therefore conclude that nuclear exclusion is the primary driver of ETV6 P214L dysfunction. Moreover, we found that the synonymous P216L mutation in mouse ETV6 (mETV6) does not create an NES due to lack of strict local sequence conservation, thus explaining the lack of hematopoietic defects in current animal models. We therefore generated *Etv6^P216L/WT^* mice with an accompanying humanizing mutation in the mutant *Etv6* allele, which are thrombocytopenic, have robust HSC defects, and serve as a new model for hematologic disease. We conclude by showing that missense mutation-dependent NESs may explain the damaging effects of other protein mutations, including POU4F3 P125L, PALB2 P378L and BRCA2 W395L. Thus, mechanistic dissection of enigmatic mutations that occur within unannotated protein regions can elucidate new modes of protein dysfunction, facilitate biomarker discovery, and inspire rational design of disease models and therapeutic strategies.

## Results

### The ETV6 P214L mutation creates an NES

The P214L mutation has no impact on recombinant ETV6 behavior or DNA binding activity in biochemical assays (Figure S1). To further assess potential P214L-dependent effects on ETV6 function, we set out to determine the minimal ETV6 construct required for P214L-dependent protein mislocalization in mammalian cells. Stable expression of full-length ETV6 constructs in HeLa cells followed by immunofluorescence analysis with an ETV6 antibody shows clear nuclear exclusion of the mutant protein (Figure 1B). Surprisingly, P214L-mediated nuclear exclusion was maintained after removal of the ETS and PNT domains (Figure S2). We note that this construct included a strong nuclear localization signal (the SV40 NLS) to compensate for loss of the endogenous ETV6 nuclear import mechanism^22^. Further truncation analysis identified residues 203-245 as the minimal ETV6 sequence required for P214L-dependent mislocalization of the mCherry-NLS construct (Figure 1C). Thus, it is clear the P214L mutation does not cause mislocalization via interference with known ETV6 activities, including DNA binding and homotypic oligomerization.

In diseases associated with an ETV6 mutation at the P214 site, patients exclusively exhibit proline substitution to a leucine residue^16,18^. To further probe the mechanism of mutation-dependent mislocalization, we interrogated the importance of residue identity at amino acid 214. Constructs bearing single base pair substitutions within the proline 214 codon were prepared and mammalian subcellular localization assays performed in the context of the mCherry-ETV6_203-245_-NLS protein (Figure 1C, S3). In all cases, including proline 214 mutation to arginine and serine, mutant protein localization was indistinguishable from wild-type ETV6. Moreover, complete removal of the proline 214 codon also yields a protein construct that localizes to the nucleus (Figure S3). Among plausible explanations, our observation that the proline 214 mutant mislocalization phenotype is specific to a leucine substitution is consistent with the creation of a protein interaction motif.

Analysis of P214L-proximal protein sequence composition revealed a preponderance of bulky aliphatic residues upstream of the mutation site, including Leu205, Ile209, and Leu212 (Figure 1D). This relatively high concentration of hydrophobic amino acids is uncommon in protein regions lacking structural order. Indeed, short leucine-rich sequences that reside within disordered protein segments were the earliest discovered ‘leucine-rich NESs’^23–25^. These NESs are directly engaged by XPO1, which is a nuclear export receptor that recognizes linear sequence motifs comprising 4-5 bulky hydrophobic residues with specific spacing patterns^26^. While the wild-type ETV6 protein has no predictable NES, the P214L mutation creates a class 1a XPO1 recognition motif from residues 205-214 (Figure 1D). Consistently, the commonly employed NES prediction software LocNES^27^ identifies a strong NES from residues 205-214 in the mutant ETV6 construct and no NES in the corresponding region of the wild-type protein (Figure S4).

### ETV6 P214L mislocalization is dependent upon a direct interaction with XPO1

To test our hypothesis that the P214L mutation causes ETV6 mislocalization via creation of an NES, we treated live HeLa cells expressing the mCherry-ETV6_203-245_-NLS construct with the small molecule XPO1 inhibitor Leptomycin B^28^ and captured fluorescent images every five minutes over a two-hour time course (Figure 1E and Supplemental Video 1). Within 30 minutes of exposure to 5 nM Leptomycin B, the mCherry-ETV6_203-245_-NLS construct had completely migrated to the nucleus in all cells observed. This experiment demonstrates that the ETV6 P214L protein is constantly being imported into the nucleus, and that ETV6 import activity is far less efficient than P214L-dependent export activity. Similar results were obtained with the ETV6 P214L construct and when Leptomycin B was replaced with the chemically distinct XPO1 inhibitor Selinexor^29^, which is an FDA-approved anti-cancer agent.

As an alternate approach to chemical XPO1 manipulation, we generated an ETV6 P214L construct with a second mutation, I209S, that abolishes the XPO1 interaction motif. The ETV6 I209S/P214L double mutant construct (henceforth ‘ETV6 P214L^nuc^’) was stably expressed in HeLa cells and imaged via immunofluorescence (Figure 1E). Consistent with chemical inhibition studies, genetic disruption of the XPO1 recognition motif fully restores nuclear localization of an ETV6 construct bearing the P214L mutation.

We next sought to validate and quantitatively assess the ETV6 P214L:XPO1 interaction in a highly controlled setting. A fluorescence polarization-based assay was devised in which a recombinantly expressed and purified complex of XPO1 and its obligate auxiliary component RanGTP^23–25,30^ was titrated into various synthetic, fluorescently-labeled ETV6_203-245_ constructs. Consistent with cellular studies, the P214L mutant construct exhibits robust binding to XPO1 whereas binding cannot be detected with the wild-type construct (Figure 1F). Notably, XPO1 engages the ETV6 P214L construct with an affinity constant (261 nM) that is comparable to that of the well-established XPO1 ligand peptide PKI (65 nM)^31^. Taken together, our results definitively demonstrate that the P214L mutation creates an XPO1 interaction motif that is responsible for ETV6 cellular mislocalization.

The heterozygous P214L mutation is known to disrupt localization of both mutant and wild-type ETV6 protein copies within the cell^16,32^, although the mechanistic basis for this observation is not known. Our data offer a rational hypothesis for the dominant negative phenotype wherein PNT domain-mediated oligomerization leads to export of mutant and wild-type ETV6 constructs. To test this, HA-tagged P214L protein and Myc-tagged wild-type protein were expressed together or separately in mammalian cells and immunofluorescence assays were performed to determine localization of each construct (Figure S5). While wild-type ETV6 on its own localizes to the nucleus as expected, co-expression with ETV6 P214L induces mislocalization of the wild-type protein to the cytoplasm. In this context, XPO1 inhibitors rescue localization of both mutant and wild-type ETV6 protein constructs. We next co-expressed wild-type ETV6 with a P214L construct bearing two PNT domain mutations (A93D and V112E) that abolish oligomerization activity^33^ (Figure S5). While the oligomerization-deficient P214L construct localizes to the cytoplasm, the wild-type construct localizes to the nucleus. Therefore, PNT domain-mediated oligomerization creates complexes of mutant and wild-type ETV6 protein copies that are exported from the nucleus by XPO1, ultimately leading to the dominant negative nature of the P214L mutation (Figure 1G).

### Restoring nuclear localization of the ETV6 P214L variant rescues transcriptional activities

We have shown that strategies to disrupt the ETV6 P214L:XPO1 interface effectively restore nuclear localization of the mutant ETV6 protein. However, it is possible that the P214L mutation also impacts ETV6 function in a manner that would render the protein inactive regardless of proper localization. To investigate this, we implemented an established cellular assay for ETV6 activity^19^. In this assay, introduction of active ETV6 suppresses growth of NRAS^G12D^-transformed Ba/F3 cells (Ba/F3^NRAS^) (Figure 2A). Wild-type, P214L, or P214L^nuc^ ETV6 constructs were stably integrated into Ba/F3^NRAS^ cells and growth kinetics were measured over a seven-day period (Figure 2B). As expected, wild-type ETV6 localizes to the nucleus and strongly inhibits NRAS^G12D^-mediated transformation while the P214L variant accumulates in the cytoplasm (Figure S6) and has no effect on Ba/F3^NRAS^ growth rate. The ETV6 P214L^nuc^ construct localizes to the nucleus and suppresses Ba/F3^NRAS^ growth in a manner that is nearly identical to that of wild-type ETV6. These results indicate that the ETV6 P214L variant can function as a suppressor of transformed Ba/F3 cell growth once nuclear localization is restored.

**Figure 2.**
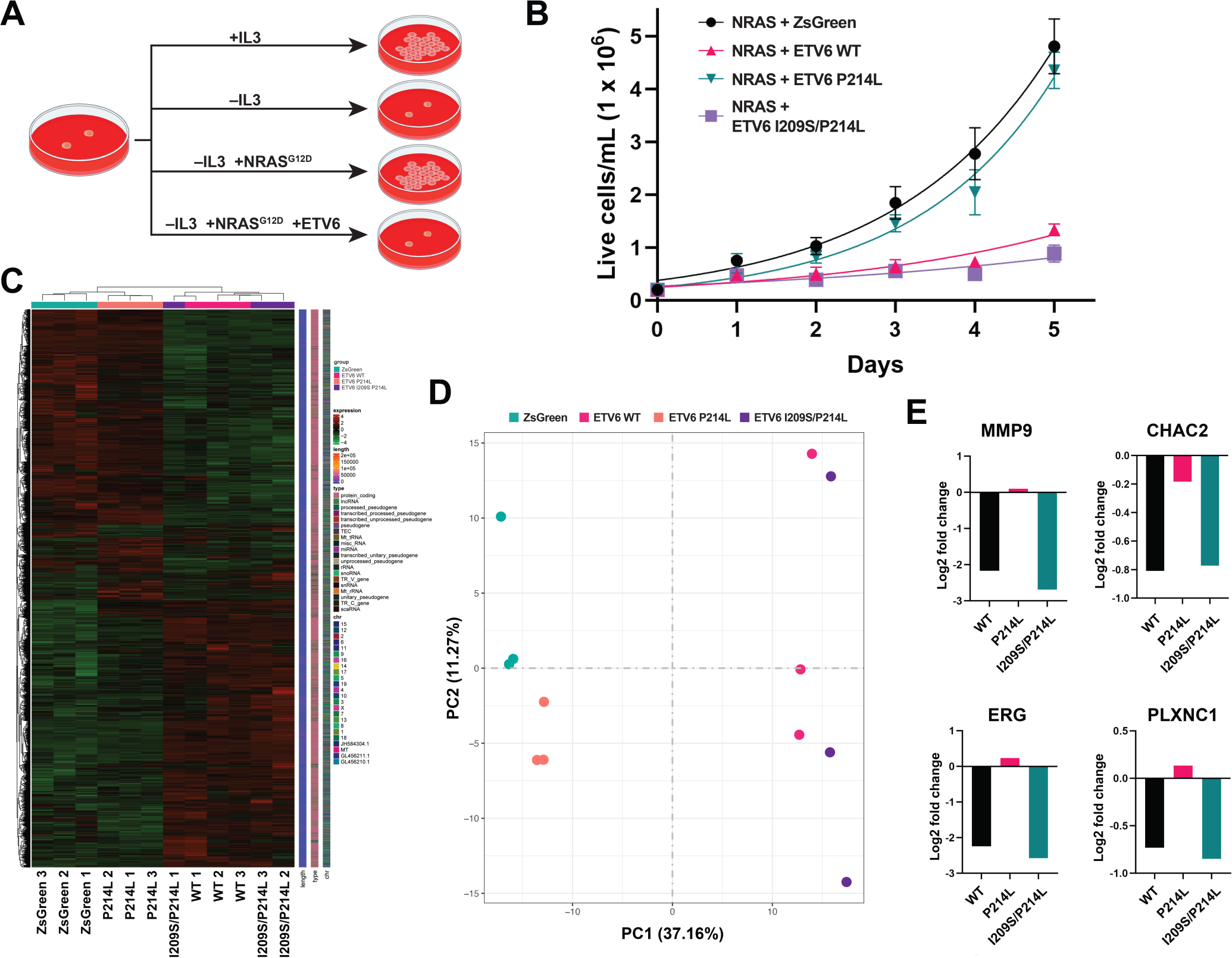
The ETV6 P214L protein exhibits wild-type ETV6 activity when nuclear localization is restored. (A) A schematic depicting the ETV6 activity in transformed Ba/F3 cells. (B) NRAS^G12D^-transformed Ba/F3 cell growth curves in the presence of ZsGreen or various ETV6 constructs. Error bars represent S.D. from n = 3 experimental replicates. (C and D) Hierarchical clustering analysis (C) and PCA analysis (D) of RNA-seq data collected from Ba/F3^NRAS^ cell lines described in (B) at the 3-day time point (n = 3). (E) Transcript abundance of known ETV6 target genes in Ba/F3^NRAS^ cell lines described in (B) at the 3-day time point (n = 3).

To further investigate the transcriptional repressor properties of the nuclear ETV6 P214L, we isolated total RNA from Ba/F3^NRAS^ cells expressing each ETV6 variant for RNA-seq characterization. Hierarchical clustering and PCA analysis of global gene expression profiles show striking overlap between Ba/F3^NRAS^ cells expressing the wild-type ETV6 and ETV6 P214L^nuc^ constructs (Figure 2C and 2D). However, very little overlap is observed between the wild-type transcriptional profile and that of the P214L variant, which closely resembles Ba/F3^NRAS^ cells lacking ETV6. We also analyzed individual transcript abundance of known ETV6 target genes, including CHAC2, ERG, MMP9, and PLXNC1 (Figure 2E). In all cases, transcript levels in wild-type and ETV6 P214L^nuc^-expressing Ba/F3^NRAS^ cells are strongly down-regulated while ETV6 P214L fails to elicit a transcriptional response. Our transcriptional profiling studies further support the conclusion that XPO1-dependent mislocalization is the primary driver of ETV6 P214L dysfunction.

### P214L-mediated phenotypes in mice require a ‘humanizing’ ETV6 mutation

Previously reported genetically engineered mice harboring a P216L mutation in the endogenous *Etv6* allele have normal platelet counts and do not exhibit an overt hematopoietic defect^21^. To investigate this perplexing observation, we introduced wild-type or P216L mETV6 constructs into NIH/3T3 cells for immunofluorescence imaging-based localization analysis (Figure 3A). Surprisingly, both constructs localize to the nucleus with equal efficiency. A sequence alignment comparing human ETV6 and mETV6 revealed that the P216L mutation does not create an XPO1 interaction motif (Figure 3B). This is due to variance at a single amino acid site, the S211 position in mice, which is an isoleucine residue in the human sequence. Coincidentally, this substitution in mETV6 precisely mimics our ETV6 P214L^nuc^ rescue construct. We found that a ‘humanizing’ mutation at the mETV6 S211 site (mETV6 S211I) does not disrupt nuclear localization while the S211I/P216L double mutation causes robust mETV6 nuclear exclusion (Figure 3A).

**Figure 3.**
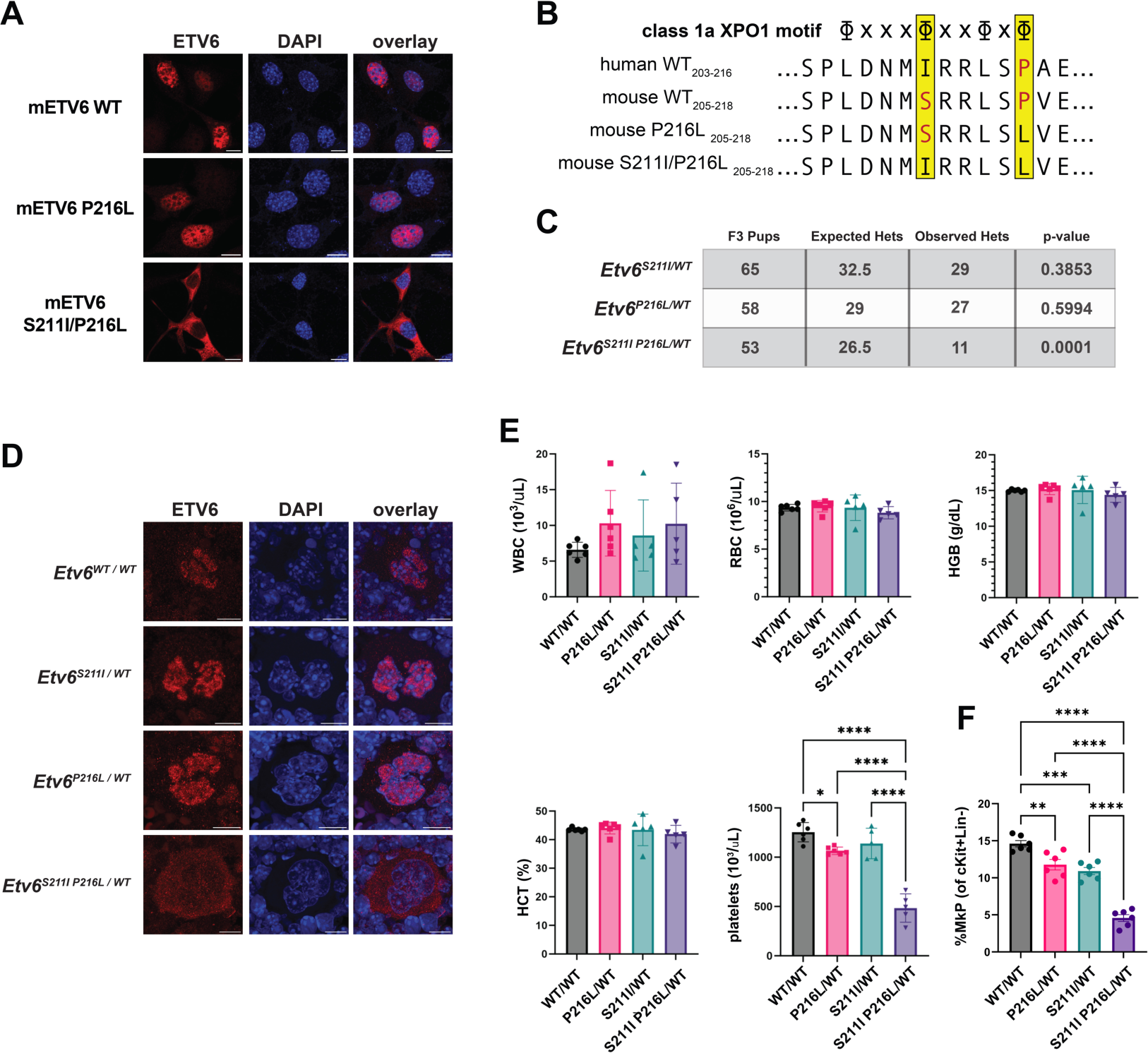
Humanization of mETV6 is required to induce P216L-dependent protein mislocalization and thrombocytopenia in mice. (A) ETV6 immunofluorescencez-stack images of NIH 3T3 cells bearing exogenous copies of the indicated full-length mETV6 construct. Scale bar = 10 μm. (B) The mETV6 protein sequence (amino acids 205-218) aligned with synonymous residues in the human ETV6 variant that comprise the ETV6 P214L-dependent XPO1 interaction motif. Red letters indicate residues that disrupts the XPO1 interaction. (C) Expected and observed heterozygous inheritance patterns for F3 generation backcross. p-value determined via Chi-squared test. (D) ETV6 immunofluorescence z-stack images of megakaryocytes in fixed mouse bone marrow from each transgenic line and wild-type littermates. Scale bar = 10 μm. (E) Complete blood counts from peripheral blood of transgenic mouse lines and wild-type littermates. Error bars represent S.E.M. from n = 5 for mETV6^S211I/WT^, n = 5 mETV6^S211I^ ^P216L/WT^, n = 6 for mETV6^WT/WT^, and n = 6 for mETV6^P216L/WT^. p-values calculated using Tukey’s multiple comparison test after one-way ANOVA. * ≤ 0.05, ** ≤ 0.01, *** ≤ 0.001, **** ≤ 0.0001. (F) Flow cytometry analysis of myeloerythroid progenitors from each genetically modified mouse. Error bars represent S.E.M. from n = 6 for each mouse genotype. p-values calculated as described in (E).

We next generated three genetically engineering mouse lines using CRISPR/Cas9 to knock-in single point mutations (S211I or P216L) or the double mutation (S211I/P216L) at the endogenous *Etv6* locus. The *Etv6^S211I/WT^*and *Etv6^P216L/WT^* single mutant mouse lines exhibited Mendelian inheritance patterns while the *Etv6^S211I^ ^P216L/WT^* double mutant line demonstrated a significant viability defect (Figure 3C). We also collected bone marrow from heterozygous mice to determine mETV6 localization in megakaryocytes via immunofluorescence (Figure 3D). Consistent with our cell culture results, mETV6 protein in *Etv6^S211I^ ^P216L/WT^* mice is mislocalized to the cytoplasm in megakaryocytes, while mice harboring a single S211I or P216L mutation exhibit proper mETV6 localization to the nucleus.

In order to determine mutation-dependent impacts on peripheral blood cell counts, CBC analysis was performed on the genetically engineered mouse lines and wild-type littermates (Figure 3E). As observed with ETV6-deficient human patients^20^, white and red blood cell counts are consistent across each genetic background. Similarly, mice carrying the mETV6 S211I or P216L mutations exhibit platelet counts resembling that of wild-type mice (*Etv6^S211I/WT^* = 1140 x 10^3^/µL ± 155.1, *Etv6^P216L/WT^* = 1065 x 10^3^/µL ± 37.82, *Etv6^WT/WT^* = 1254 x 10^3^/µL ± 98.61). However, all *Etv6^S211IP216L/WT^* double mutant mice are thrombocytopenic with platelet counts ∼60% lower than wild-type littermates (485 x 10^3^/µL ± 143.8). Notably, human *ETV6^P214L/WT^* patients exhibit strikingly similar platelet deficiencies, with platelet count decreases of ∼70% compared to non-carrier relatives^20^. Additional characterization of myeloerythroid progenitors^34^ revealed a decrease in megakaryocyte progenitors in *Etv6^S211I^ ^P216L/WT^* mice (Figure 3F and S7), phenocopying the megakaryocyte maturation defect that has been described in patients^16^. Thus, as our biochemical studies suggest, the humanizing S211I mutation in mice is essential to properly model human ETV6 P214L dysfunction. Given the clear thrombocytopenia phenotype, the *Etv6^S211I^ ^P216L/WT^* mouse may also prove useful in the study of additional ETV6 mutation-associated blood diseases, including MDS and leukemia.

### The humanized mETV6 P216L mutation causes hematopoietic stem cell functional defects

Beyond phenocopying the megakaryocyte maturation and thrombocytopenia that characterizes patients with ETV6 mutations, the *Etv6^S211I^ ^P216L/WT^* mouse also represents a new *in vivo* system to study the impact that mutation-engendered ETV6 dysfunction has on HSCs. Previous work has shown that complete loss of *Etv6* is embryonic lethal and that conditional inactivation of *Etv6* in the hematopoietic system leads to depletion of HSCs in adult mouse bone marrow^4,5^. Given that *Etv6^S211I^ ^P216L/WT^*mice are viable, in contrast to the phenotype of the *Etv6* knockout mouse, we wondered if these mice would exhibit HSC defects. Flow cytometry analysis of bone marrow hematopoietic stem and progenitor populations revealed a significant decrease in the absolute number of Lin^−^Sca-1^−^c-Kit^+^(L^−^S^−^K^+^) cells and HSCs in our *Etv6^S211I^ ^P216L/WT^* mice compared to *Etv6^WT/WT^* littermates (Figures 4A and S8). Notably, there was no difference in the frequency of myeloid progenitor populations, suggesting that mETV6 S211I/P216L may specifically impact HSCs.

**Figure 4.**
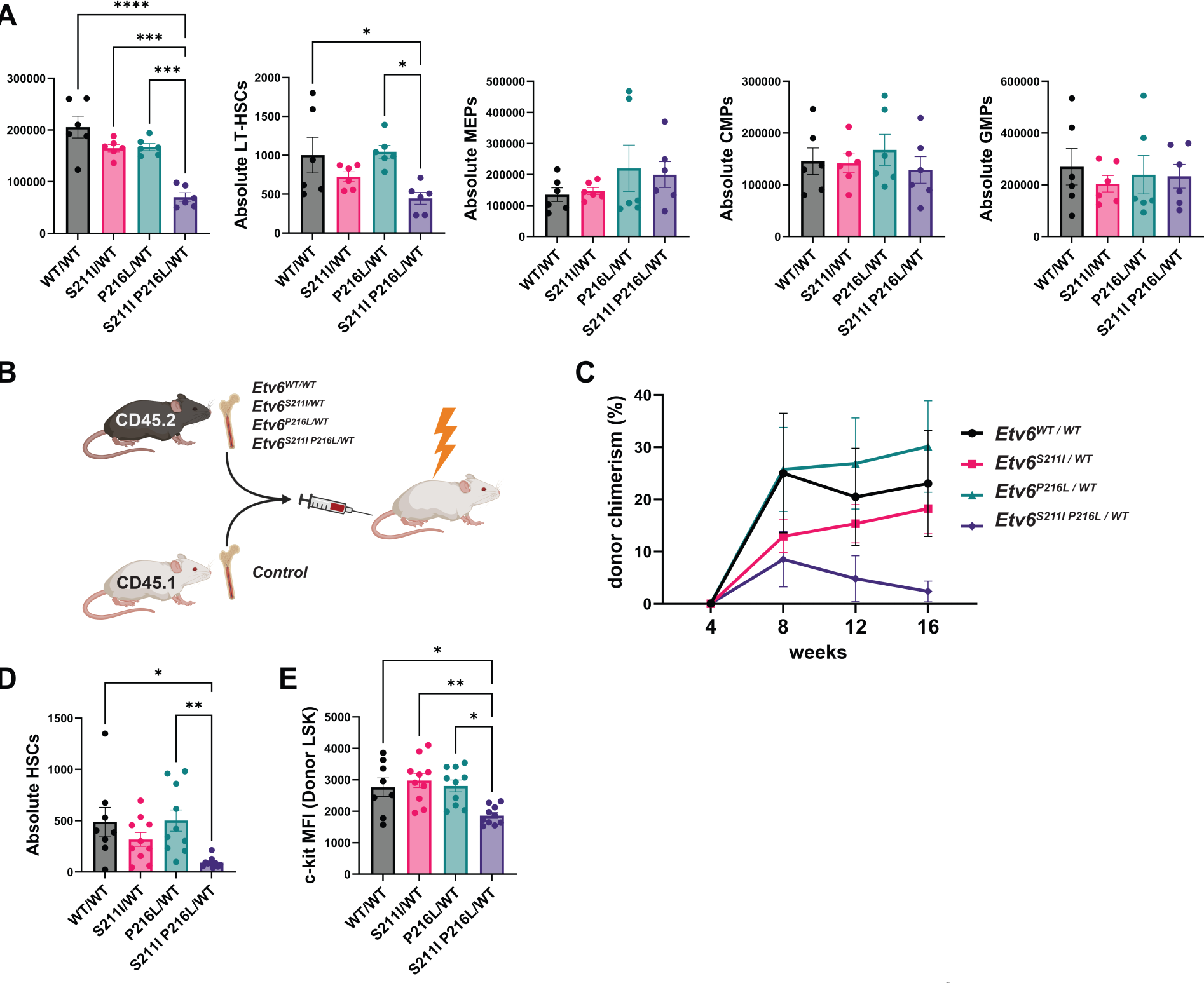
The *Etv6^S211I^ ^P216L^* allele causes hematopoietic stem cell defects in mice. (A) Flow cytometry analysis of bone marrow cell populations from wild-type and ETV6 mutant mice. Error bars represent S.E.M. from n = 6 mice. p-values calculated using Tukey’s multiple comparison test after one-way ANOVA. * ≤ 0.05, ** ≤ 0.01, *** ≤ 0.001, **** ≤ 0.0001. (B) Competitive bone marrow transplant experimental design. (C) Competitive bone marrow transplant engraftment results. Donor chimerism represents the percent of peripheral blood cells harboring the CD45.2 antigen as measured via flow cytometry. Error bars represent S.E.M. for n = 8 for test bone marrow from mETV6^WT/WT^, n = 10 for test bone marrow from mETV6^S211I/WT^, n = 10 for test bone marrow from mETV6^P216/WT^, and n = 10 for test bone marrow from mETV6^S211I^ ^P216L/WT^. (D) Competitive bone marrow transplant endpoint analysis of absolute CD45.2 positive HSCs as measured via flow cytometry. Error bars represent S.E.M for n = 6 mice from each genotype. p-values calculated as in (A). (E) Competitive bone marrow transplant endpoint analysis of c-kit mean fluorescence intensity in CD45.2 positive LSK cells as measured via flow cytometry. p-values calculated as in (A).

To test whether mETV6 S211I/P216L leads to a cell autonomous defect in HSC self-renewal, we performed competitive bone marrow transplant assays wherein equal numbers of bone marrow cells from control mice and ETV6 mutant mouse lines were co-injected into lethally irradiated recipient mice and peripheral blood donor engraftment was measured over a 16-week period (Figure 4B). We found that *Etv6^S211I^ ^P216L/WT^-*derived bone marrow cells exhibited a defect in engraftment in recipient mice when compared to mice that received cells from *Etv6^S211I/WT^*, *Etv6^P216L/WT^*, and *Etv6^WT/WT^* littermates (Figure 4C). After 16 weeks, mice were sacrificed, and hematopoietic stem and progenitor cell populations were analyzed. Of the genotypes tested, only donor *Etv6^S211I^ ^P216L/WT^* HSCs were significantly decreased in number and self-renewal quotient compared with donor HSCs from wild-type littermates, with the average donor chimerism value dropping to ∼2.4% (Figure 4D). Additionally, in donor *Etv6^S211I^ ^P216L/WT^* cells there was a decrease in the frequency of HSCs exhibiting high levels of c-Kit expression, a population that exhibits an intrinsic megakaryocyte lineage bias (Figure 4E)^35^. Collectively, our HSC analyses demonstrate that while P214L-mediated ETV6 dysfunction does not cause strict embryonic lethality, the mutation does have profound impacts on HSC maintenance and function. The *Etv6^S211I^ ^P216L/WT^* HSC defects may also explain the high incidence rate of MDS and leukemia in human *ETV6^P214L/WT^*carriers^16,18,20^, as discussed in detail below.

### Missense mutations create NESs in multiple disease-associated proteins

Mutations that disrupt protein folding to unmask an otherwise inaccessible NES or induce a frameshift that creates an NES have been previously reported^36–39^. The ETV6 P214L missense mutation is unique in that it directly converts the surrounding sequence into a nuclear export signal. Unlike the previously reported mechanisms, we were able to envision a straightforward bioinformatic search to identify missense mutations that directly create an NES (Figure 5A). To perform such a search, we first identified all mutations in the ClinVar database that are classified as “Pathogenic”, “Likely Pathogenic”, or “Variant of Unknown Significance” and reside within regions of nuclear proteins lacking predictable structure elements, as defined by AlphaFold. We next input all wild-type and corresponding mutant protein sequences into the NES prediction program LocNES and identified mutations that: (i) create a strong NES (LocNES score > 0.4), and (ii) have a LocNES score >0.3 points higher than that of the wild-type counterpart sequence. For these proof-of-concept screens, we focused on X-to-L mutations only. We identified 73 unique protein mutations, including ETV6 P214L, that match the above criteria and create an NES (Table 1 and Supplemental Dataset 1).

**Figure 5.**
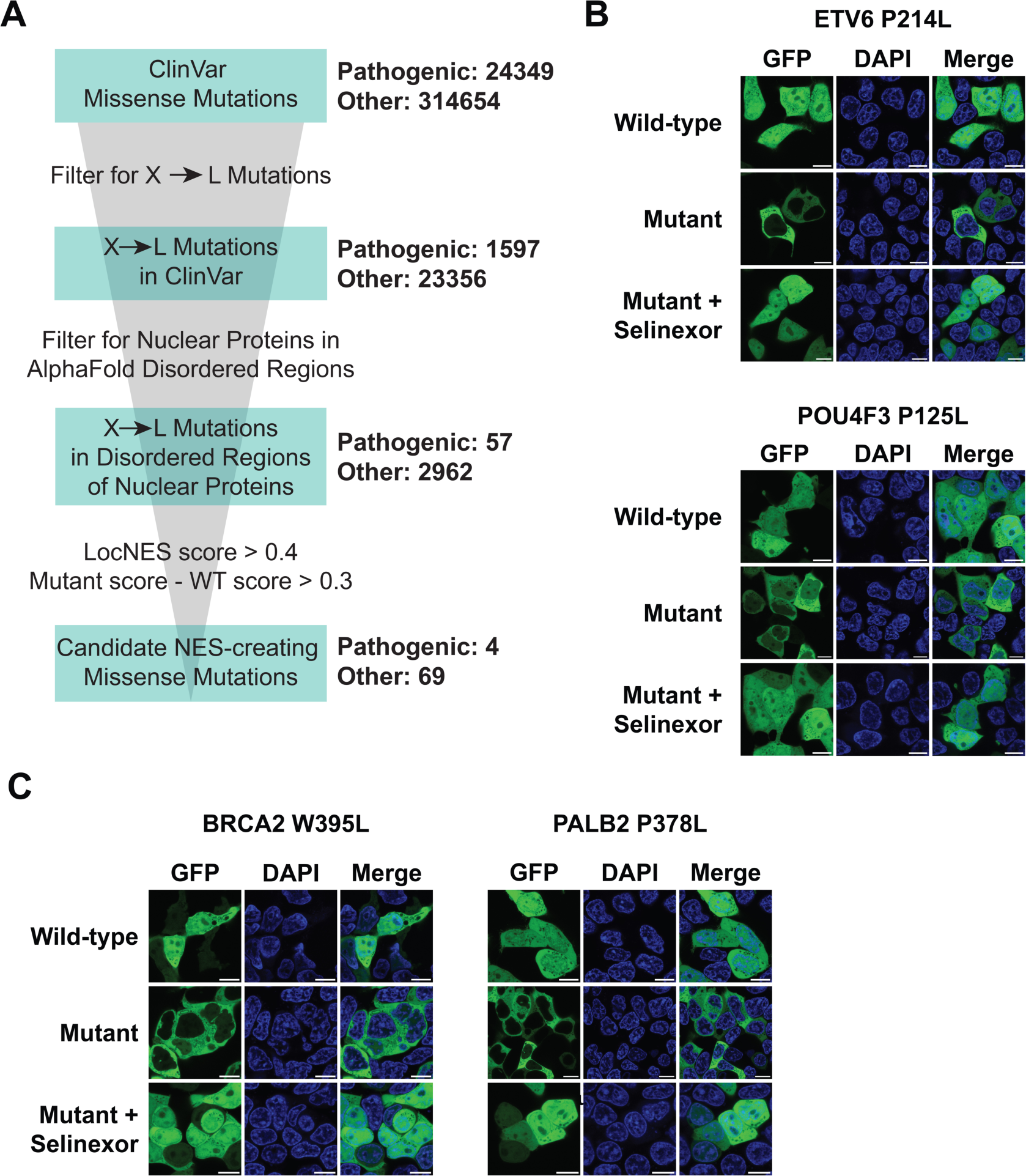
A bioinformatics screen identifies mutation-dependent NESs in diverse nuclear proteins and diseases. (A) Summary of bioinformatics screen to identify disease-associated mutations that create candidate NESs. (B) Fluorescence confocal microscopy images of live HEK293T cells bearing GFP-fused ETV6 fragments (amino acids 203-233) and POU4F3 fragments (amino acids 105-135) from mutant proteins and wild-type counterparts. Images post-Selinexor treatment for 2 h are indicated. Scale bar = 10 μm. (C) As in (B) with GFP-fused BRCA2 fragments (amino acids 380-410) and PALB2 fragments (amino acids 364-394) from mutant proteins and wild-type counterparts.

**Table 1.** Top candidate NES-creating mutations in the ClinVar database.

| Pathogenic NES Candidate Mutations |  |  |  |  |
| --- | --- | --- | --- | --- |
| Gene | Mutation | Mutant LocNES | WT LocNES | ΔLocNES |
| ETV6 | P214L | 0.611 | 0 | 0.611 |
| BCOR | P85L | 0.494 | 0 | 0.494 |
| POU4F3 | P125L | 0.621 | 0.153 | 0.468 |
| FOXP3 | P75L | 0.546 | 0.241 | 0.305 |
| Top 10 Likely Pathogenic/Variant of Unknown Significance NES Candidate Mutations |  |  |  |  |
| Gene | Mutation | Mutant LocNES | WT LocNES | ΔLocNES |
| ACD | Pro256Leu | 0.795 | 0 | 0.795 |
| SLX4 | Ser1157Leu | 0.764 | 0 | 0.764 |
| PANK2 | Arg44Leu | 0.723 | 0 | 0.723 |
| ZIC1 | Pro144Leu | 0.715 | 0 | 0.715 |
| ANKRD11 | Pro1848Leu | 0.684 | 0 | 0.684 |
| MEF2C | Pro175Leu | 0.656 | 0 | 0.656 |
| FOXP3 | Pro133Leu | 0.644 | 0 | 0.644 |
| QRICH1 | Pro284Leu | 0.623 | 0 | 0.623 |
| PRX | Pro1309Leu | 0.685 | 0.101 | 0.584 |
| MEFV | Ser273Leu | 0.583 | 0 | 0.583 |
| Homology-directed Repair Protein NES Candidate Mutations |  |  |  |  |
| Gene | Mutation | Mutant LocNES | WT LocNES | ΔLocNES |
| BRCA2 | His1362Leu | 0.78 | 0 | 0.78 |
| BRCA1 | Pro1091Leu | 0.691 | 0 | 0.691 |
| BRCA1 | Pro115Leu | 0.665 | 0 | 0.665 |
| BRCA1 | Pro724Leu | 0.663 | 0 | 0.663 |
| BRCA2 | Ser636Leu | 0.583 | 0 | 0.583 |
| PALB2 | Pro378Leu | 0.583 | 0 | 0.583 |
| BRCA1 | Pro633Leu | 0.525 | 0 | 0.525 |
| BRCA2 | Trp395Leu | 0.519 | 0 | 0.519 |
| BRCA1 | Pro1603Leu | 0.527 | 0.051 | 0.476 |
| BARD1 | Trp218Leu | 0.476 | 0 | 0.476 |

Excluding the ETV6 P214L mutation, three hits from our screen are characterized as ‘pathogenic’ in the ClinVar database (BCOR P85L, POU4F3 P125L, and FOXP3 P75L). None of these mutations have an established mechanism of protein dysfunction. Interestingly, the FOXP3 mutation occurs within a previously characterized NES and so was not pursued further here^40^. Fluorescently labeled peptides corresponding to BCOR P85L and POU4F3 P125L were synthesized and employed as ligands in our fluorescence polarization-based XPO1 interaction assay (Figure S9). While BCOR did not interact with XPO1 in this assay, POU4F3 P125L showed a robust, mutation-dependent interaction with reconstituted export machinery. We next turned to a mammalian cell NES characterization assay wherein the putative NES is tethered to the C-terminus of an otherwise diffuse cellular GFP construct. Upon confirmation of assay efficacy with the ETV6 P214L mutation (Figure 5B), we characterized localization of POU4F3 mutant and wild-type constructs. Consistent with cellular NES activity, the POU4F3 P125L peptide blocks nuclear localization of GFP while the wild-type counterpart has no impact on protein localization (Figure 5B). Furthermore, the mutation-dependent mislocalization phenotype can be rescued via treatment with Selinexor. We therefore propose that the POU4F3 P125L causes non-syndromic autosomal recessive deafness^41^ through the XPO1-dependent mislocalization mechanism. Notably, POU4F3 is a transcription factor that is essential for proper inner ear hair cell differentiation, and loss-of-function mutations are known to cause deafness.

We next employed the cellular GFP-NES characterization assay to interrogate: (i) the top-ranked likely pathogenic/variants of unknown significance from our screen, and (ii) mutations targeting well-established genes in homology-repair (HR)-deficient cancers. The latter set was included to identify new biomarkers for HR-deficient cancers, for which treatment strategies such as PARP inhibitors are available and may be immediately useful^42^. Five variants of unknown significance, including two HR-related mutations (ACD P256L, ANKRD11 P1848L, MEF2C P175L, BRCA2 W395L and PALB3 P378L), exhibit robust mutation-and XPO1-dependent mislocalization (Figure 5B and S10). We therefore propose that these mutations are genetic markers for protein dysfunction. Moreover, our findings demonstrate the efficacy of computational methods to identify mutation-dependent NESs.

## Discussion

Here we have found that the enigmatic ETV6 P214L mutation creates an XPO1-dependent NES, and that nuclear export activity is required for mutation-associated hematopoietic defects. This unexpected discovery underscores the value of mechanistic dissection as it pertains to mutations that occur within unannotated protein regions. Mechanistic insight enabled rational design of genetic and small molecule approaches to rescue ETV6 P214L localization in cells and clearly explains the dominant negative phenotype. We were also able to address the perplexing observation that the currently studied mETV6 P216L mouse fails to recapitulate most human disease phenotypes. Indeed, design of the humanized mETV6 mice described here, which properly model P214L-dependent dysfunction in hematopoiesis, was inspired by our mechanistic studies. Beyond ETV6, the mutation-dependent export pathway reported herein can be extrapolated to other disease-associated protein mutations of unknown function. Considering that our search was confined to X-to-L mutations in the ClinVar database and only the most well-characterized NES motifs, we expect more exhaustive searches will continue to fuel discovery of new genetic biomarkers for protein dysregulation. Notably, this mechanism is distinct from and potentially more damaging than traditional loss-of-function mutations in that it can also cause mislocalization of wild-type protein copies or other nuclear proteins by forming complexes that are shuttled out of the nucleus.

The previously reported *Etv6^P216L/WT^* and *Etv6^P216L/P216L^* mice exhibit only subtle hematopoietic deficits in very specific contexts^21^, raising questions about the discrepancies between mouse and human phenotypes. Our humanized *Etv6^S211I^ ^P216L/WT^*mouse represents the first genetically accurate model of ETV6 thrombocytopenia that recapitulates the degree of thrombocytopenia and megakaryocyte progenitor defects observed in patients. Additionally, we uncover a role for ETV6 P216L in impairing HSC self-renewal, and we propose that mutation-dependent HSC defects observed in mice explain why human carriers are strongly predisposed to MDS and leukemia. Of note, ETV6 has been demonstrated to repress inflammatory response genes^43^, and elevated innate inflammation is a hallmark of MDS stem cells^44^. Future studies are needed to determine whether ETV6 P216L is sufficient for transformation to MDS with aging, or whether cooperating mutations, as have been described in patients^18^, are required. Our work provides a much-needed model to carry out such studies. Finally, we observe in humanized *Etv6^S211I^ ^P216L/WT^* mice depletion of HSCs with high expression of c-Kit, a population previously described to have a megakaryocyte lineage bias^35^, with megakaryocytes arising directly from such HSCs^45^. This raises the possibility that the megakaryocyte progenitor defect and thrombocytopenia associated with ETV6 mutations arise directly from a defect in HSCs, which would mirror a similar mechanism described for RUNX1 mutation-associated thrombocytopenia^46^.

The missense mutation-dependent NES prediction workflow described here was effective in identifying mutant sequences that engage XPO1. However, there are many factors that mitigate prediction reliability. The efficiency of an endogenous NLS relative to the corresponding mutation-dependent NES can impact protein nucleocytoplasmic distribution. Current NLS and NES prediction tools must integrate several variables, including numerous motifs, disorder propensity, and are likely missing yet-to-be-discovered signals^27^. These variables temper the reliability and accuracy of predictions. Additionally, prediction software cannot account for: (i) specific import and export receptor activities in a given cell type, (ii) NLS/NES accessibility in the cellular environment, or (iii) post-translational modifications that regulate translocation. Thus, experimental validation is currently a necessary part of our workflow to validate mutation-dependent NESs. Moving forward, efforts to better understand the rules and regulators of nucleocytoplasmic transport are necessary to enable higher throughput identification of mutation-dependent protein mislocalization events in disease.

The class of NES-creating mutations defined in our work may be immediately combatable with existing therapeutic regimes. Our results indicate that small molecule XPO1 inhibitors, including the FDA-approved chemotherapeutic Selinexor, may be successfully repurposed to rescue nuclear localization of proteins bearing a mutation-dependent NES. We also note that by clarifying variants of unknown significance, our study and follow-up efforts may elucidate new genetic biomarkers for disease predispositions and effective treatment strategies. For example, patients carrying the BRCA2 W395L or PALB2 P378L mutations should be considered strong candidates for chemotherapeutic agents that target HR-deficient cancers. Given the implications from our study, we believe unannotated protein regions and associated genetic mutations offer a fertile starting point for biological discovery.

## Supplemental information

Supplemental Information can be found online at xxx.

**Supplemental Video 1**. Time-lapse fluorescence confocal microscopy analysis of live HEK293T cells expressing mCherry-ETV6 P214L_203-245_-NLS-GFP. Cells were treated with 5 nM leptomycin B at t = 0 and z-stack images were captured every 6 min for a total of 84 min. Scale bar = 10 μm.

**Supplemental Dataset 1.** Results from the bioinformatics search of the ClinVar database to identify X-to-L mutations that create candidate NESs. Tabs for ‘Pathogenic’ and ‘Likely Pathogenic/VUS’ are indicated.

## Acknowledgements

We thank the Ben Tu Lab for assistance with confocal microscopy data collection, the UT Southwestern Transgenic Core Facility for assistance with transgenic mice, the Sean Morrison Lab for assistance with CBC data collection, the UT Southwestern Tissue Management Shared Resource for assistance with bone marrow immunofluorescence staining, and the Histo Pathology Core for assistance with bone marrow embedding and sectioning. This work was supported by grants from the Cancer Prevention and Research Institute of Texas (RR180051 to G.L.), the Welch Foundation (I-2039-20230405 to G.L. and I-1532 to Y.M.C.), the American Heart Association (937595 to G.L.), and the NIH (5F30HL167629 to M.M., 1R35GM147140 to G.L., and R35GM141461 to Y.M.C.). G.L. is a Virginia Murchison Linthicum Scholar in Medical Research. Y.M.C. is the Alfred and Mabel Gilman Chair in Molecular Pharmacology and the Eugene McDermott Scholar in Biomedical Research (Y.M.C.).

## Author contributions

G.L., M.M., and R.B conceived the study. G.L., M.M., R.B., S.C., and T.T. designed experiments. M.M., T.T., R.B., C.V., S.C., Y.C., and G.L. contributed to data curation and formal analysis. G.L., M.M., S.C., and Y.C. secured funding for the work. G.L. and M.M. wrote the original draft with input from S.C. All authors reviewed and edited the final version of the manuscript.

## STAR Methods

### Key resources table

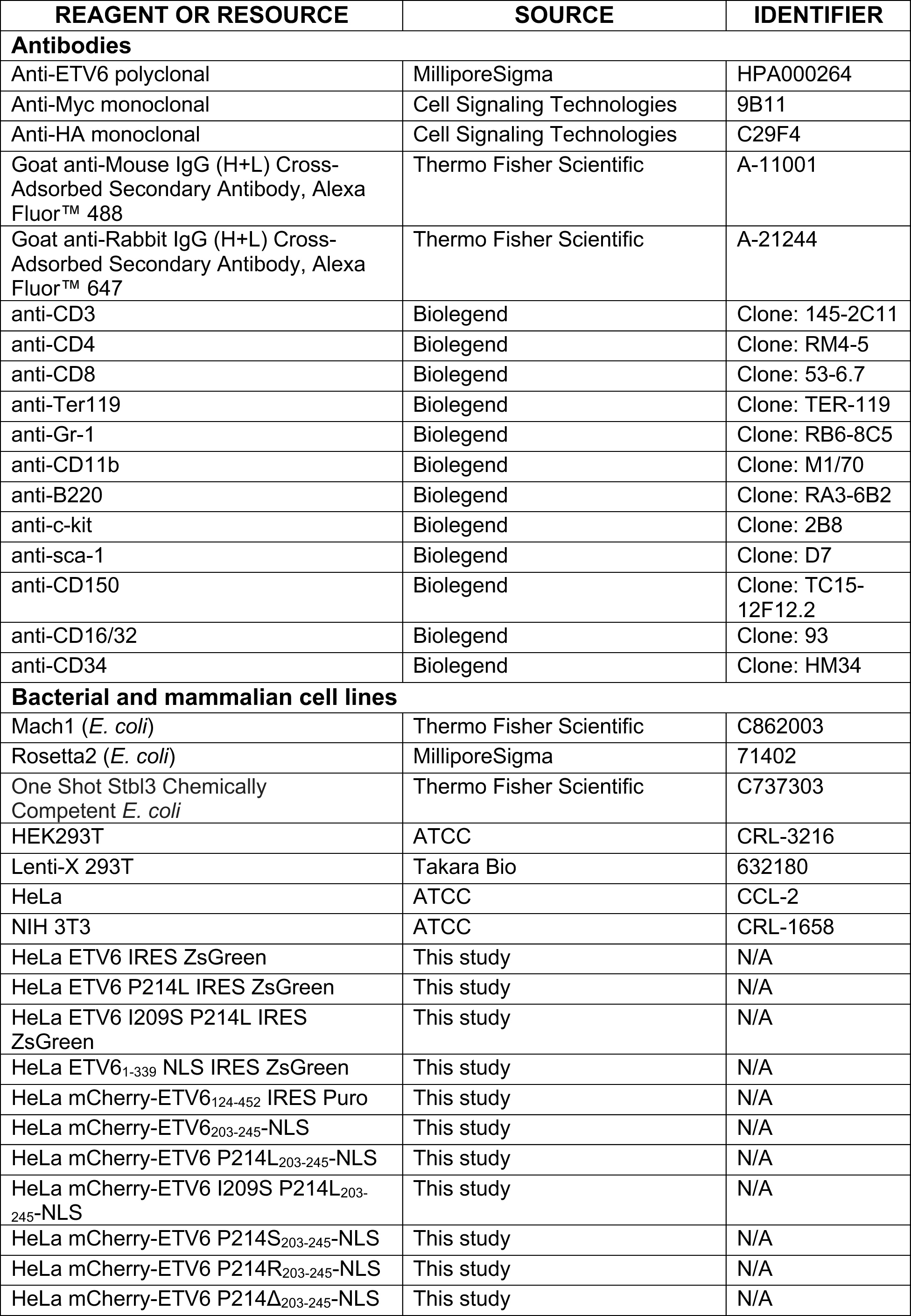
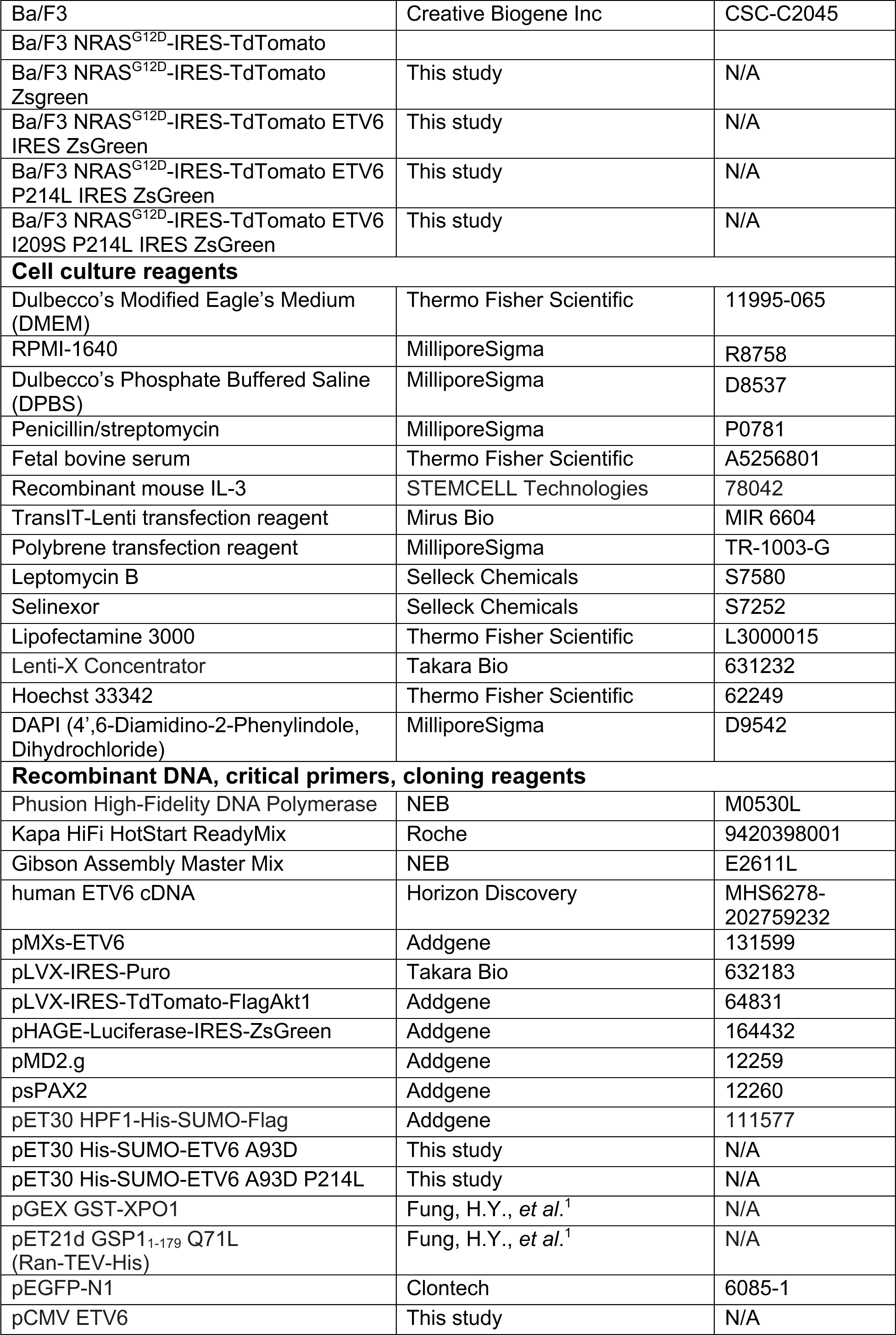
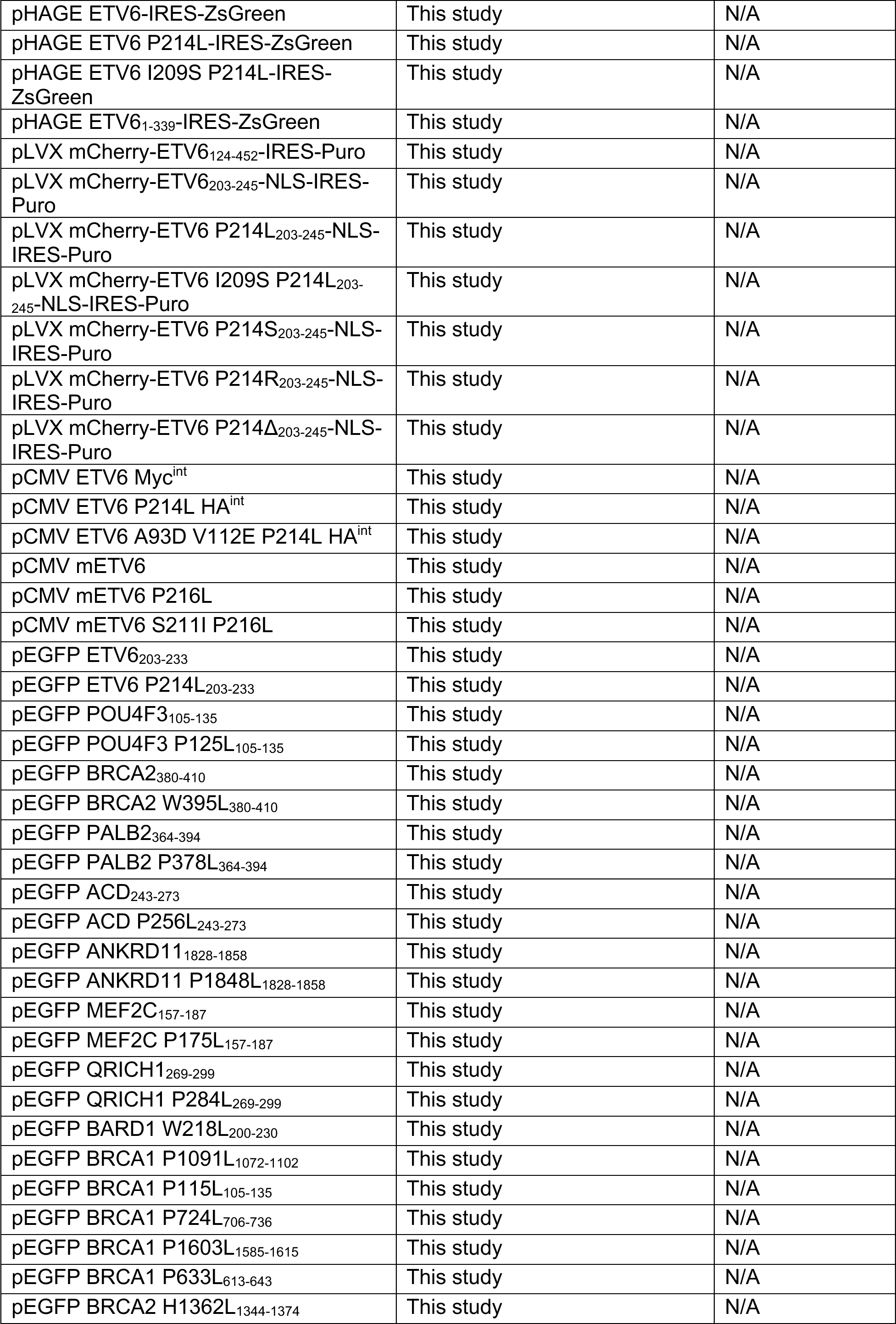
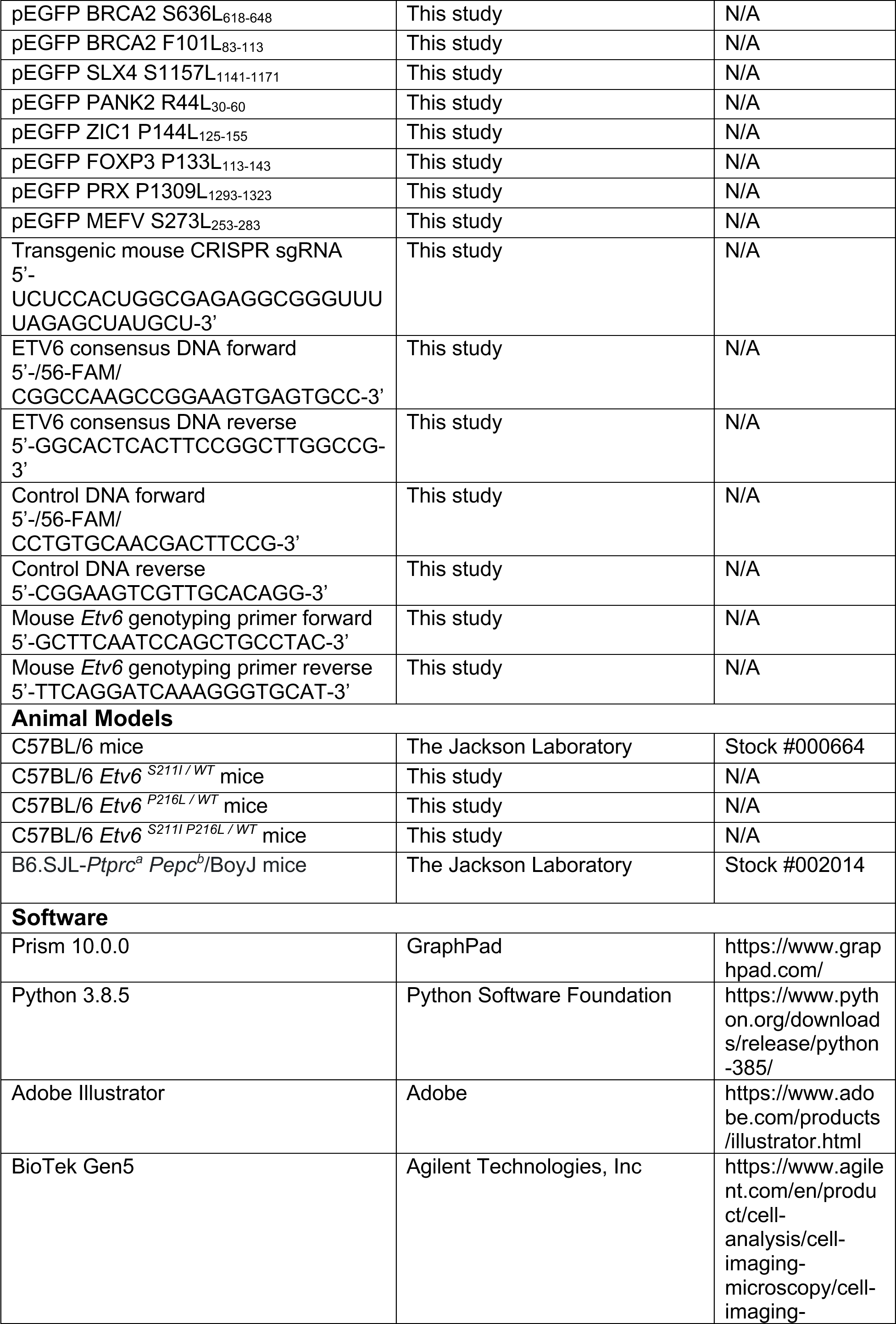
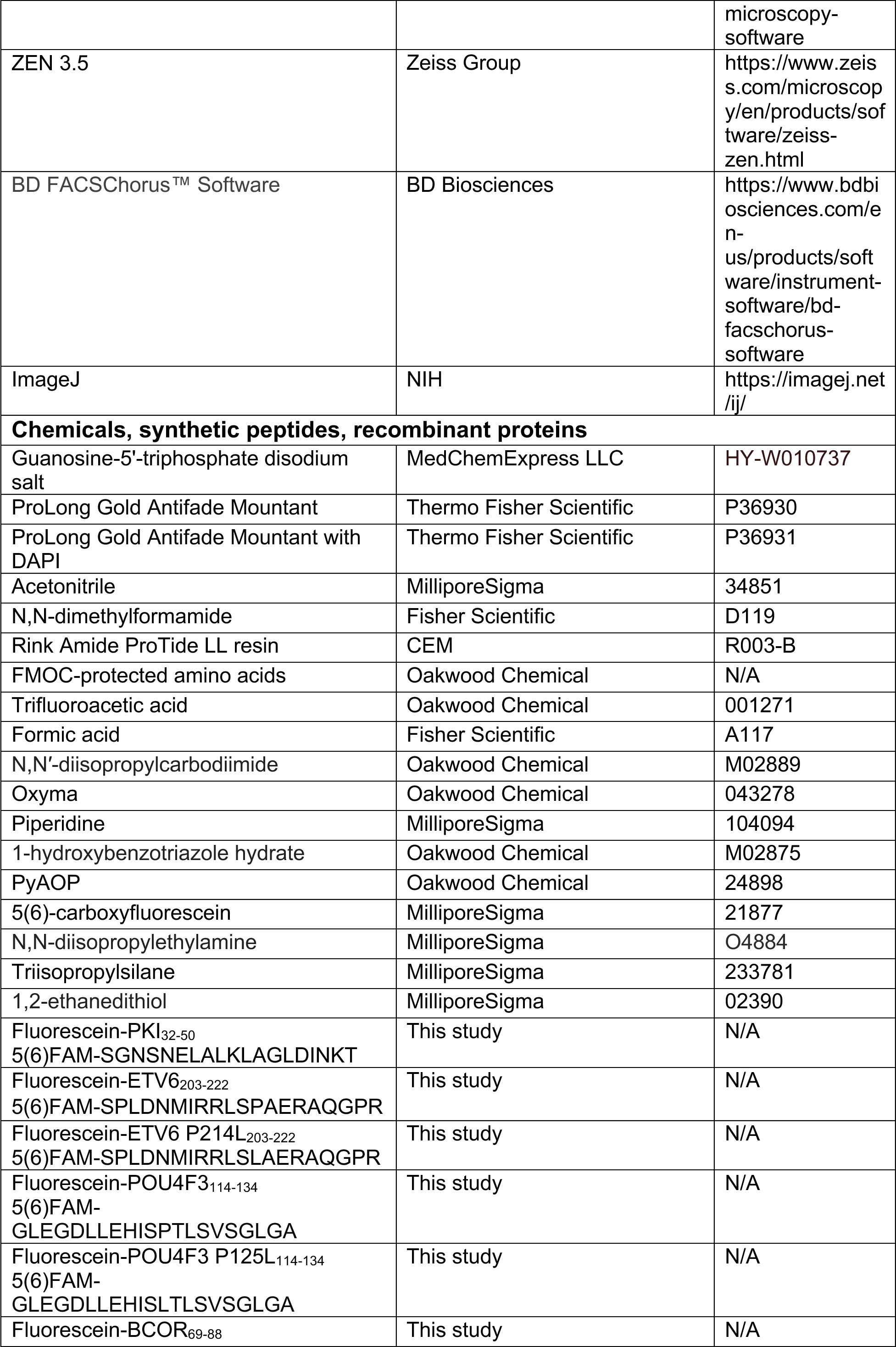
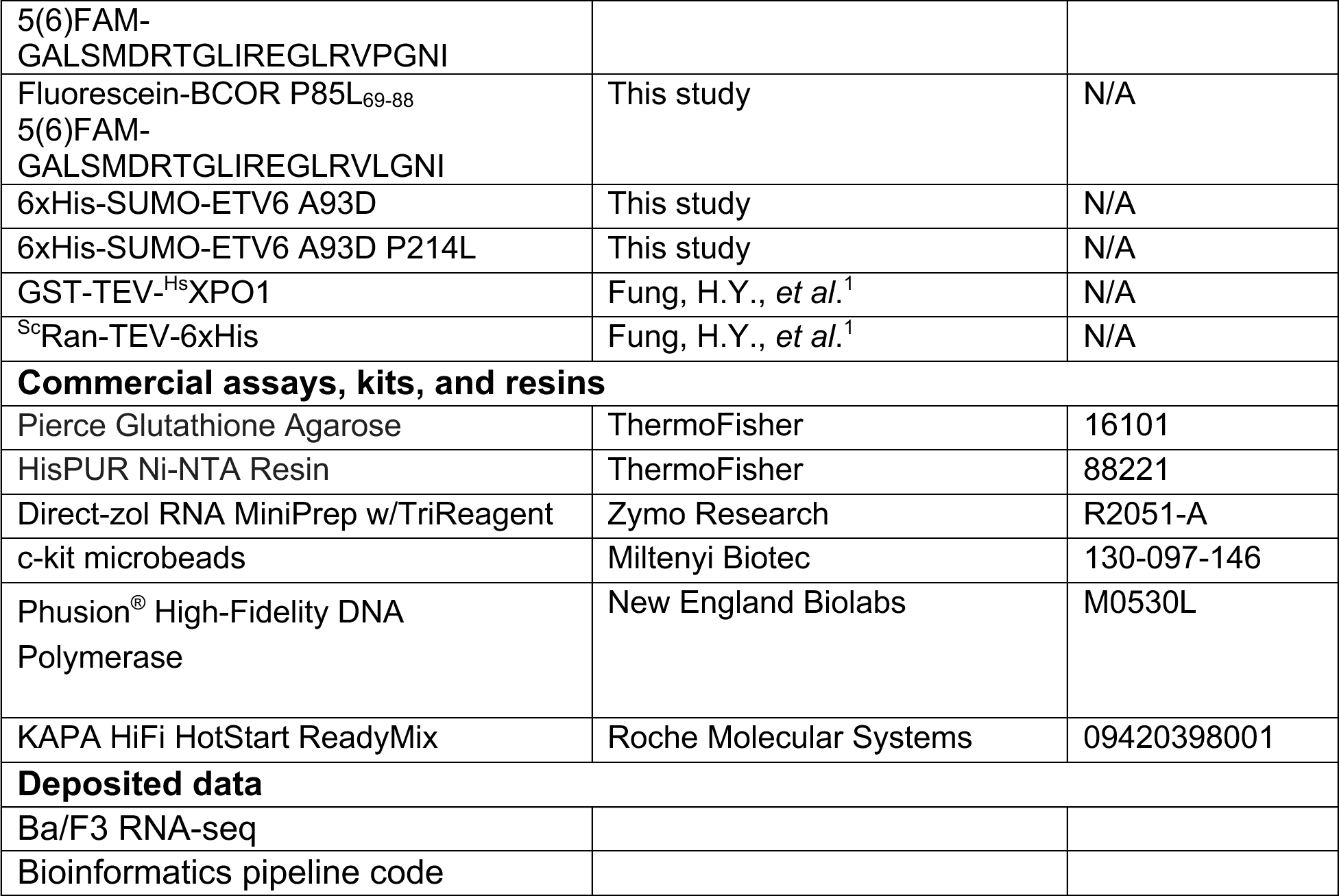

### Molecular cloning

All PCR amplification steps were performed using the Phusion High-Fidelity DNA Polymerase or KAPA HiFi DNA Polymerase according to the Manufacturer’s Protocols. Point mutations were installed via inverse PCR. All DNA oligonucleotides were synthesized by Sigma-Aldrich (Milwaukee, WI) or Integrated DNA Technologies (Coralville, IA). All plasmids used in this study were sequence verified by GENEWIZ (South Plainfield, NJ) or EurofinsGenomics (Louisville, KY). Lentivirus plasmid preparation was performed using restriction enzyme cloning and plasmid amplification was carried out in Stbl3 *E. coli* cells. All other cloning was carried out using Mach1 *E. coli* cells. To install mutations into lentivirus plasmids, mutant genes were amplified from pET30, pCMV, or pEGFP transient mammalian expression vectors and subcloned into the desired lentivirus vector via restriction enzyme cloning.

### Recombinant protein expression and purification

pET30 plasmids encoding 6xHis-SUMO-ETV6 A93D or 6xHis-SUMO-ETV6 A93D P214L protein were transformed into Rosetta2 (DE3) cells and inoculated into 6 L of Luria Broth (Miller). Cells were grown in a shaker at 37°C to an OD_600_ of 0.6 and protein expression was induced with 0.5 mM IPTG at 18°C overnight. Cells were sonicated in lysis buffer containing 50 mM Tris, pH 7.5, 500 mM NaCl, 5 mM β-ME and 1 mM PMSF. Soluble lysate was isolated via centrifugation at 48,000 RCF for 40 min at 4°C. The soluble fraction was incubated with Ni-NTA resin for 1 h at 4°C and the resin was subsequently washed with 50 column volumes of lysis buffer supplemented with 25 mM imidazole. After washing, the protein was eluted in lysis buffer supplemented with 300 mM imidazole. The elution was dialyzed into a buffer containing 50 mM Tris, pH 7.5, 500 mM NaCl, and 2 mM TCEP for 16 h at 4°C in the presence of the Ulp1 protease to cleave the 6xHis-SUMO tag. The dialysate was then concentrated to 2 mL using an Amicon Ultra Centrifugal filter (Millipore; 30 kDa MWCO) and injected onto a gel filtration column (HiLoad 16/600 Superdex 200; Cytiva) that had been pre-equilibrated with a buffer containing 50 mM Tris, pH 7.5, 150 mM NaCl, 10% glycerol, and 2 mM TCEP. Pure fractions (as judged by SDS-PAGE) were concentrated, flash frozen in single-use aliquots, and stored at –80°C.

Human XPO1 *^(Hs^*XPO1) and yeast Ran (*^Sc^*Ran) were purified according to previously published methods^1^. Briefly, a pGEX-TEV vector encoding *^Hs^*XPO1 and a pET21d vector encoding the yeast homolog of Ran (GSP1 residues 1–179, Q71L; gift from Dr. Takuya Yoshizawa) were transformed separately into Rosetta2 (DE3) cells and inoculated into 6 L of Luria Broth (Miller). Cells were grown in a shaker at 37 °C up to an OD_600_ of 0.6 for *^Sc^*Ran and 0.8 for *^Hs^*XPO1. *^Sc^*Ran protein expression was induced with 0.5 mM IPTG at 18°C for 16 h and *^Hs^*XPO1 protein expression was induced with 0.5 mM IPTG at 25°C for 10 h.

Cells expressing *^Hs^*XPO1 were lysed by sonication in buffer containing 40 mM HEPES (pH 7.5), 2 mM MgOAc, 200 mM NaCl, 10 mM dithiothreitol (DTT), 10% glycerol, and protease inhibitors. Soluble lysate was isolated via centrifugation at 48,000 RCF for 40 min at 4°C. The soluble fraction was incubated with Pierce glutathione agarose (Thermo Fisher Scientific) for 1 h at 4°C. The agarose was washed with 50 column volumes of lysis buffer. *^Hs^*XPO1 was then cleaved with TEV protease for 1 h at 4°C, further purified by size-exclusion chromatography (20 mM HEPES pH 7.5, 100 mM NaCl, 5 mM Mg_2_OAc, and 2 mM TCEP), flash frozen in single-use aliquots, and stored at –80°C.

Cells expressing *^Sc^*Ran were lysed in buffer containing 50 mM Tris (pH 7.5), 2 mM Mg_2_OAc, 500 mM NaCl, 2 mM DTT, and protease inhibitors. *^Sc^*Ran was then purified by affinity chromatography with Ni-NTA resin as above with 6xHis-SUMO-ETV6 constructs. After affinity purification, *^Sc^*Ran was loaded with GTP (incubated with molar excess of ethylenediaminetetraacetic acid (EDTA) for 30 min on ice followed by incubation with excess GTP and Mg_2_OAC for 30 min at room temperature) and then purified by ion exchange chromatography (HiTrap SP, Cytiva). *^Sc^*Ran was further purified by size-exclusion chromatography (20 mM HEPES pH 7.5, 110 mM K_2_OAc, 2 mM Mg_2_OAc, 10% glycerol, and 2 mM TCEP), flash frozen in single-use aliquots, and stored at –80°C.

### Fluorescence polarization-based DNA-binding assay

Oligos for fluorescence polarization assays were ordered from Integrated DNA Technologies (IDT). The sequences for oligos with and without ETV6 consensus sequences (in bold) are below:

ETV6 consensus DNA forward: /56-FAM/CGGCCAAGCC**GGAA**GTGAGTGCC

ETV6 consensus DNA reverse: GGCACTCAC**TTCC**GGCTTGGCCG

Control DNA forward: /56-FAM/CCTGTGCAACGACTTCCG

Control DNA reverse: CGGAAGTCGTTGCACAGG

ETV6 A93D or ETV6 A93D/P214L at varying concentrations were incubated for 20 min at 4°C with 5 nM ETV6 DNA or Control DNA. Proteins and DNA were diluted using a buffer containing 50 mM Tris, pH 7.5, 50 mM NaCl, 2 mM MgCl_2_, 0.01% Triton X-100, and 2 mM DTT. 50 μL reactions were added to a black, flat-bottom 96-well plate (Corning Costar) and analyzed on a BioTek Cytation 5 imager equipped with a Green FP filter set (excitation: 485 nm, emission: 528 nm). Polarization in the absence of titrant was background subtracted from all data points.

### Peptide synthesis

#### General protocols

All fluorenylmethyloxycarbonyl (Fmoc)-protected amino acids were purchased from Oakwood Chemical or Combi-Blocks. Rink Amide ProTide LL peptide synthesis resin was purchased from CEM. All analytical reversed-phase HPLC (RP-HPLC) was performed on an Agilent 1260 series instrument equipped with a quaternary pump and an XBridge Peptide C18 column (5 μm, 4 × 150 mm; Waters) at a flow rate of 1 mL/min. Preparative RP-HPLC was performed on an Agilent 1,260 series instrument equipped with a preparatory pump and a XBridge Peptide C18 preparatory column (10 µM; 19 × 250 mm, Waters) at a flow rate of 20 mL/min. All instruments were equipped with a variable wavelength UV-detector. All RP–HPLC steps were performed using 0.1% (trifluoroacetic acid, TFA) in H_2_O (Solvent A) and 90% acetonitrile, 0.1% TFA in H_2_O (Solvent B) as mobile phases. For LC/MS analysis, 0.1% formic acid (Sigma-Aldrich) was substituted for TFA in mobile phases. Mass analysis was carried out for each product on an LC/MSD (Agilent Technologies) equipped with a 300 SB-C18 column (3.5 µM; 4.6 × 100 mm, Agilent Technologies).

Peptide identities (N-terminal fluorescein, C-terminal amidation)

PKI_32-50_: 5(6)FAM-SGNSNELALKLAGLDINKT

ETV6_203-222_: 5(6)FAM-SPLDNMIRRLSPAERAQGPR

ETV6 P214L_203-222_: 5(6)FAM-SPLDNMIRRLSLAERAQGPR

POU4F3_114-134_: 5(6)FAM-GLEGDLLEHISPTLSVSGLGA

POU4F3 P125L_114-134_: 5(6)FAM-GLEGDLLEHISLTLSVSGLGA

BCOR_69-88_: 5(6)FAM-GALSMDRTGLIREGLRVPGNI

BCOR P85L_69-88_: 5(6)FAM-GALSMDRTGLIREGLRVLGNI

#### Synthetic procedures

The above amidated peptides were synthesized via solid-phase peptide synthesis on a CEM Discover Microwave Peptide Synthesizer (Matthews, NC) using the Fmoc-protection strategy on Rink Amide-ChemMatrix resin (0.5 mmol/g). For coupling reactions, amino acids (5 eq) were activated with N,N′-diisopropylcarbodiimide (DIC, 5 eq)/Oxyma (5 eq) and heated to 90°C for 2 min while bubbling with nitrogen gas in N,N-dimethylformamide (DMF). Fmoc deprotection was carried out with 20% piperidine in DMF supplemented with 0.1 M 1-hydroxybenzotriazole hydrate (HOBt) at 90°C for 1 min while bubbling with nitrogen gas.

To add an N-terminal fluorescein label, 5(6)-carboxyfluorescein (3 eq) was activated with PyAOP (3 eq) and N,N-diisopropylethylamine (DIPEA, 6 eq) and coupled to the deprotected α-amine on resin for 30 min at 25°C in DMF while bubbling with nitrogen gas. Resin was washed with DMF and treated with 20% piperidine in DMF. Peptide cleavage from the resin was performed with 92.5% TFA, 2.5% triisopropylsilane (TIS), 2.5% 1,2-ethanedithiol (EDT), and 2.5% H_2_O for 2 h at 25°C. The crude peptide was then precipitated by the addition of a 10-fold volume of cold ether and centrifuged at 4000 RCF for 10 min at 4°C. The pellet was resuspended in Solvent A and purified via preparative RP-HPLC using a linear gradient from 0 to 70% Solvent B over 30 min. Fractions were analyzed on analytical RP-HPLC and ESI-MS and those containing pure product (>95%) were pooled, lyophilized, and stored at –80°C.

### Fluorescence polarization-based peptide-binding assay

Varying concentrations of XPO1:RanGTP (1:3 ratio of XPO1 to RanGTP) were incubated for 20 min at 4°C with 5 nM fluorescently labeled peptide. XPO1:RanGTP and peptides were diluted using a buffer containing 20 mM HEPES, pH 7.5, 100 mM NaCl, 2 mM Mg_2_OAc, 0.01% Triton X-100, 2 mM DTT, and 10% glycerol. 50 µL reactions were added to a black, flat-bottom 96-well plate (Corning Costar) and analyzed on a BioTek Cytation 5 imager equipped with a Green FP filter set (excitation: 485 nm; emission: 528 nm). Polarization in the absence of titrant was background subtracted from all data points.

### Cell culture

HEK293T cells, HeLa cells, and NIH/3T3 cells were cultured in DMEM supplemented with 10% Heat-Inactivated FBS and penicillin/streptomycin at 37°C and 5% CO_2_. Ba/F3 cells were cultured in RPMI 1640 supplemented with 10% Heat-Inactivated FBS, penicillin/streptomycin, and 10 ng/mL recombinant mouse IL-3 at 37°C and 5% CO_2_.

### Generation of Lentivirus for stable ETV6 overexpression cell lines

All lentivirus production was done using the HEK293T Lenti-X cell line and the pMD2.g and psPAX2 lentivirus packaging plasmids. To stably express ETV6 constructs for localization studies, all constructs were PCR amplified and cloned into either the pLVX-IRES-Puro vector using the EcoRI and BamHI restriction sites or the pHAGE-Luciferase-IRES-ZsGreen vector using the NotI and BamHI restriction sites.

On day 1, Lenti-X cells were seeded into 6-well plates. On day 2, each well was transfected with 1 µg pLVX or pHAGE expression plasmid, 0.6 µg psPAX2 and 0.4 µg pMD2.g packaging plasmids, and 4 µL TransIT-Lenti transfection reagent. On day 3, FBS was supplemented to 30%. On day 4, virus-containing media in each well was collected and replaced with DMEM supplemented to 30% FBS. On day 5, virus-containing media was again collected and pooled with media collected the previous day. All supernatant was syringe filtered using 0.22 µm PES filters (Cytiva). This filtered virus was concentrated using the Lenti-X Concentrator (Takara Bio Inc) and then stored at −80°C.

### Stable expression and localization of ETV6 constructs

To stably express ETV6 constructs in HeLa cells, cells were seeded into a 10 cm plate in media supplemented with 8 µg/mL polybrene. While HeLa cells were still in suspension, lentivirus was added to the media. 48 h later, 2 µg/mL puromycin was added to the media to select for cells infected with pLVX-IRES-Puro or cells were sorted for green fluorescence using a BD Melody fluorescence-activated cell sorter for cells infected with pHAGE-Luciferase-IRES-ZsGreen based virus.

After selecting for populations stably expressing the desired constructs, HeLa cells were seeded onto round glass coverslips in 24 well plates. The following day, the cells were washed with phosphate buffered saline (PBS) 3 times and fixed with 10% formaldehyde in PBS for 15 min. Cells expressing an mCherry-tagged construct were then stained with DAPI (0.1 µg/mL) and washed 3 times before mounting on glass microscope slides with ProLong Gold Antifade Mountant. Cells expressing constructs requiring immunofluorescence for localization analysis were again washed 3 times with PBS, followed by permeabilization and blocking with a buffer containing 5% normal goat serum and 0.3% Triton-X 100 in PBS (blocking buffer) for 1 h. Cells were then washed 3 times with PBS and incubated with anti-ETV6 primary antibody diluted 1:250 in a buffer containing 1% normal goat serum and 0.1% Tween20 (staining buffer) overnight at 4°C. The following day, cells were washed 3 times with PBS and incubated with anti-rabbit IgG secondary antibody conjugated to Alexa Fluor 647 diluted 1:500 in staining buffer for 1 hr at 4°C. Cells were then stained with DAPI (0.1 µg/mL) and washed 3 times before mounting on glass microscope slides with ProLong Gold Antifade Mountant.

### Dominant negative localization assay

To separately visualize wild-type ETV6 and ETV6 P214L within the same cells, we placed an internal Myc tag in the disordered linker of the pCMV ETV6 plasmid (between amino acids 132 and 133) and an HA tag at the same locus of the pCMV ETV6 P214L plasmid using inverse cloning. We then added 2 missense mutations, A93D and V112E, in the PNT domain of the pCMV ETV6 P214L HA^int^ plasmid to render the expressed protein incapable of oligomerization^2^.

HEK293T cells were seeded onto round glass coverslips pre-treated with poly-D-lysine in 24 well plates. pCMV ETV6 Myc^int^ and pCMV ETV6 P214L HA^int^ plasmids were transfected into HEK293T cells individually to confirm localization was unchanged. pCMV ETV6 Myc^int^ was then co-transfected with pCMV ETV6 P214L HA^int^ or pCMV ETV6 A93D V112E P214L HA^int^. Cells were allowed 24 h to express protein. Additionally, to test the ability of XPO1 inhibitors to rescue localization of both wild-type and mutant protein, cells co-expressing pCMV ETV6 Myc^int^ and pCMV ETV6 P214L HA^int^ were treated with 1 µM Selinexor for 2 h. All cells were washed, fixed, permeabilized, and blocked as above for cells stably expressing human ETV6 constructs. Cells were incubated with anti-Myc and anti-HA primary antibodies (Cell Signaling Technologies) diluted 1:1000 in staining buffer overnight at 4°C. The following day, cells were washed 3 times with PBS and incubated with anti-mouse IgG secondary antibody conjugated to Alexa Fluor 488 and anti-rabbit IgG secondary antibody conjugated to AlexaFluor 647 diluted 1:500 in staining buffer for 1 hr at 4°C. Cells were the stained with DAPI (0.1 µg/mL) and washed 3 times before mounting on glass microscope slides with ProLong Gold Antifade Mountant.

### Transient mETV6 overexpression and localization

The murine *Etv6* gene was subcloned from the pMXs-ETV6 plasmid into our pCMV-ETV6 vector using Gibson Assembly, giving pCMV-mETV6. Inverse cloning was used to produce the S211I and P216L mutations in the pCMV-mETV6 plasmid to create pCMV-mETV6 P216L and pCMV-mETV6 S211I P216L.

NIH 3T3 cells were seeded onto round glass coverslips in 24 well plates. The following day, cells were transfected with mETV6 plasmids using Lipofectamine 3000 (Thermo Fisher Scientific). After 24 hours of expression, cells were washed, fixed, permeabilized, blocked, stained, and mounted as above for cells stably expressing human ETV6 constructs.

### Confocal Imaging

Mounted cells from above experimental procedures were all imaged using a Zeiss LSM 980 equipped with either a Plan-Apochromat 63x/1.40 oil objective or a Plan-Apochromat 100x/1.40 oil objective. Images were further processed in ImageJ with the Fiji image processing package.

Images were cropped and brightness adjusted in ImageJ to allow for optimal visualization of protein localization. Z-stacked images were transformed using max intensity projection.

### Time-lapse ETV6 P214L localization rescue with XPO1 inhibitor

To prevent degradation of the disordered linker and NLS in our pCMV mCherry-ETV6 P214L_203-245_-NLS, we added GFP to the C-terminus of the construct using Gibson Assembly to give pCMV mCherry-ETV6 P214L_203-245_-NLS-GFP. HEK293T cells were transfected with this construct in 35 mm glass bottom dishes. After 24 hours of expression, 5 nM Leptomycin B was supplemented to the media and time lapse imaging of selected cells began. Images were captured every 6 min for 84 min using a Zeiss LSM 880 with a Plan-Apochromat 63x/1.4 oil objective. Images were cropped and brightness adjusted in ImageJ to allow for optimal visualization of protein localization. Z-stacked images were transformed using max intensity projection.

### Ba/F3 Growth Curves

To stably express the constitutively active NRAS^G12D^ in Ba/F3 cells, a gene block corresponding to the NRAS^G12D^ gene was ordered from Integrated DNA Technologies and assembled into the pLVX-IRES-TdTomato-FlagAkt1 plasmid using the EcoRI and BamHI restriction sites. Lentivirus containing this construct was made as described above and virus was added to the Ba/F3 cells in suspension along with 8 ng/mL polybrene. Cells were centrifuged at 1,000 x g for 1 h at 25°C to increase transduction efficiency. A population of positively transduced Ba/F3 NRAS^G12D^-IRES-TdTomato cells was sorted using a BD Biosciences FACSMelody Cell Sorter.

To stably express ZsGreen or ETV6 constructs in Ba/F3 NRAS^G12D^-IRES-TdTomato cells, cells were transduced with lentivirus containing ETV6 constructs in the same manner as described above for NRAS^G12D^-IRES-TdTomato. At 48 h post-transduction, cells were sorted using a BD Melody fluorescence-activated cell sorter to select for cells stably expressing both TdTomato and ZsGreen. DAPI (1 µg/mL) was used to select for live cells, and a population of untransduced Ba/F3 cells were used as negative controls to establish gates for TdTomato and ZsGreen. Cells were sorted into 6 well plates in RPMI 1640 + 10% FBS + penicillin/streptomycin without IL-3 at a density of 2 x 10^5^ cells/mL. Media was refreshed every 48 h. Cells were counted daily or every other day using Trypan Blue staining with a BioRad TC20 Cell Counter to generate growth curves.

### Ba/F3 RNA sequencing

Ba/F3 cells were transduced and sorted as above. RNA was isolated from Ba/F3 cells 72 h after sorting into IL-3 deficient media using the Zymogen Direct-zol RNA Miniprep Kit. Isolated RNA was frozen and stored at −80°C. RNA sample quality control, mRNA library preparation, next generation sequencing, and quantification analysis was performed by Novogene Co, LTD (Beijing, China). Briefly, sample quantitation, integrity, and purity quality control was performed using the Agilent 5400 Fragment Analyzer System. mRNA library construction with poly A enrichment was followed by quality assessment with Qubit and real-time PCR for quantification, and bioanalyzer for size distribution detection. PE150 reads were sequenced on the Illumina NovaSeq 6000 platform. Raw fastq files were mapped to the mm39 reference genome using HISAT2^3^. Quantification of mapped reads was performed using featureCounts^4^ and DESeq2^5^ was used for differential expression analysis. Genes with adjusted p-value <= 0.05 and |log2(FoldChange)| >= 0 were considered as differentially expressed.

### Ba/F3 ETV6 localization

Ba/F3 cells were transduced with pHAGE based ETV6 lentivirus in the same manner as described above. At 48 h post-transduction, cells were sorted using a BD Melody fluorescence-activated cell sorter to select for cells stably expressing both ZsGreen. DAPI (1 µg/mL) was used to select for live cells, and a population of untransduced Ba/F3 cells were used as negative controls to establish the ZsGreen gate. Cells were sorted into 24 well plates containing poly-D-lysine coated round glass coverslips. The cells were centrifuged at 1,000 x g for 30 min at 25°C. Cells were then washed, fixed, permeabilized, blocked, stained, and mounted as above for cells stably expressing human ETV6 constructs. Cells were imaged using a Zeiss LSM 980 equipped with a Plan-Apochromat 63x/1.40 and further processed in ImageJ with Fiji image processing package. Images were cropped and brightness adjusted in ImageJ to allow for optimal visualization of protein localization.

### Mice

All mouse experiments were approved by the University of Texas Southwestern Medical Center Institutional Animal Care and Use Committee (IACUC) and performed in accordance with National Institutes of Health guidelines (animal protocol numbers: 2022-103291 and 2019-102633).

### Generation of genetically engineered mice

*Etv6^S211I/WT^, Etv6^P216L/WT^, and Etv6^S211I^ ^P216L/WT^* mice were generated using CRISPR/Cas9-mediated genome editing at the University of Texas Southwestern Transgenic Core Facility. Oligos harboring the desired point mutations were co-injected with Cas9–sgRNA complexes into the pronucleus of fertilized C57BL/6 eggs. Founders carrying the desired alleles were maintained in a pure C57BL/6J background with continuous backcrossing. Mice were genotyped using Sanger sequencing.

### Peripheral blood collection for CBC Analysis

Tail vein blood collections were performed on F3 generation male mice at 6 weeks of age. Blood was collected by dripping directly into K_3_ EDTA coated blood collection tubes (Greiner Bio-One MiniCollect). CBC analysis was performed using an Element 5HT Veterinary Hematology Analyzer (Heska Corporation) according to standard Manufacturer’s Instructions.

### Bone marrow immunofluorescence staining and confocal microscopy

Femurs were dissected from adult male mice and a razor blade was used to cut along the edge of the bone to expose the marrow. The marrow was then fixed in 10% Neutral Buffered Formalin for 72 h. Fixed bone marrow was removed from the surrounding bone, embedded in paraffin blocks, and sectioned for staining.

Immunofluorescence staining was performed on a Leica Bond RX. Briefly, the slides were baked for 30 min in the oven at 60°C, then the slides were loaded into the Leica Bond RX to deparaffinize and hydrate before the antigen retrieval step. Heat-induced antigen retrieval was performed at pH 6 for 20 min at 100°C in a Leica Bond RX. The slides were incubated with primary antibody anti-ETV6 (Sigma, HPA000264) (1:200 dilution) for 20 min followed by the secondary antibody goat anti-rabbit IgG secondary Alexa Fluor 647 (Invitrogen) (1:500 dilution) for 30 min. All slides were mounted with ProLong™ Gold Antifade Mountant with DNA Stain DAPI (Invitrogen, Cat P36931).

Cells were imaged using a Zeiss LSM 980 equipped with a Plan-Apochromat 63x/1.40 and further processed in ImageJ with Fiji image processing package. Images were cropped and brightness adjusted in ImageJ to allow for optimal visualization of protein localization. Z-stack images were transformed with max intensity projection.

### Flow cytometry and cell sorting of mouse bone marrow

Flow cytometry and fluorescence-activated cell sorting was performed on a BD Biosciences FACSAria II Cell Sorter. Analysis and isolation of mouse hematopoietic cells was performed using the following antibodies: CD3 (145-2C11), CD4 (RM4-5), CD8 (53-6.7), Ter119 (TER-119), Gr-1 (RB6-8C5), CD11b (M1/70), B220 (RA3-6B2), c-kit (2B8), sca-1 (D7), CD150 (TC15-12F12.2), CD16/32 (93), CD34 (HM34)from Biolegend. A lineage stain consisting of CD3, CD4, CD8, Ter119, Gr-1, CD11b, and B220 was used. Dead cells were excluded from analyses and cell sorting using propidium iodide (0.1 μg/ml). Cells stained with CD34 antibodies were incubated for 90 min on ice, with all other incubations for 30 min on ice.

### Bone marrow cell isolation

Bone marrow (BM) cells were isolated for analysis by flushing two femurs of adult male mice. For HSC-isolations, cells were isolated by crushing the spine, femurs, tibias, and pelvic bones of mice with a mortar and pestle into phosphate-buffered saline (PBS) with 2% fetal bovine serum, followed by filtration through a 70 μm strainer. C-kit enrichment was completed using c-kit microbeads (Miltenyi Biotec) and following the standard Manufacturer’s Protocol.

### Competitive transplantation assay

Female C57BL/6 mice (CD45.1 Pep boy, The Jackson Laboratory) were used as recipient mice in transplant experiments. Recipient mice were lethally irradiated with two 550 cGy doses (XRAD 320 irradiator, Precision X-Ray) given at least 3 h apart. Within 24 h of irradiation, 5.0 x 10^5^ donor cells and 5.0 x 10^5^ recipient cells were transplanted via retro-orbital injection.

### Peripheral blood chimerism analysis

Blood was collected by lateral tail vein collection using capillary blood collection tubes (Greiner Bio-One MiniCollect). Donor peripheral blood (PB) chimerism was measured in recipient mice every four weeks. PB specimens underwent red blood cell lysis with ammonium chloride potassium (ACK) lysis buffer (Gibco) followed by staining with antibodies against CD45.1, CD45.2, Gr-1, CD11b, B220, and CD3. Complete blood counts were obtained using a Procyte Dx Hematology Analyzer (IDEXX) according to standard Manufacturer’s Instructions.

### Endpoint bone marrow analysis

At 16 weeks after transplant, BM cells were isolated from two femurs of recipient mice as detailed above and stained with antibodies against CD45.1, CD45.2, lineage (CD3, CD4, CD8, B220, CD11b, Gr-1, Ter119), c-kit, sca-1, CD48, and CD150).

### Bioinformatics scan for additional NES-creating missense mutations

The dataset of ClinVar mutations used in our analysis was downloaded on 4/17/2021 from https://ftp.ncbi.nlm.nih.gov/pub/clinvar/tab_delimited/variant_summary.txt.gz. Predicted structures of the *H. sapiens* proteome were downloaded from AlphaFold (Reference Proteome UP000005640). Bioinformatics pipeline was written using Python 3.8.5 utilizing the Biopython package.

The ClinVar dataset was filtered for mutations resulting in a change from any amino acid to leucine. The “X-to-L” dataset was then further filtered for mutations occurring in disordered regions of nuclear proteins. A “disordered region” in a given protein was defined as any 25 amino acid stretch in which each residue had an AlphaFold pLLDT score <= 0.65. A list of proteins annotated as nuclear (GO:0005634) was downloaded from Uniprot and used to filter for nuclear proteins.

The filtered X-to-L dataset was used to generate a list of mutant protein sequences that were bulk uploaded to the online LocNES prediction tool alongside corresponding wild-type sequences. Candidate *de novo* NESs were defined as having a mutant LocNES score > 0.4 at a position lacking any NES in the wild-type sequence or a mutant LocNES score at least 0.3 greater than the wild-type LocNES score at the same position.

### Localization of GFP-tagged candidate NESs

Codon optimized gene blocks encoding the mutant sequences from the top hits in the bioinformatics scan were ordered from IDT. Gibson assembly was used to insert these 31 amino acid sequences on the C-terminus of GFP, creating GFP-31mer plasmids. HEK293T were transfected with GFP-tagged mutant candidate NES sequences using Lipofectamine 3000 (Thermo Fisher Scientific Inc.) in 35 mm glass bottom dishes (Cellvis). 24 hours after transfection, the nuclei were stained with 10 µM Hoechst 33342 in PBS for 10 min. Cells were washed 3 times with complete media and imaged by a Zeiss LSM 980 equipped with a Plan-Apochromat 100x/1.40 oil objective. Mutant sequences that resulted in decreased nuclear GFP signal were again transfected and imaged after treatment with 1 µM Selinexor for 2 h. Additionally, corresponding wild-type sequences were created using inverse cloning. Wild-type sequences transfected and imaged in the same manner as the mutant sequences. Images were cropped and brightness adjusted using ImageJ and Fiji image processing package for optimal visualization of GFP localization.

### Statistical analysis

All statistical analysis was done using GraphPad Prism 10.0.0.

