## Supplemental Figures for "Enigmatic missense mutations can cause disease via creation of *de novo* nuclear export signals"

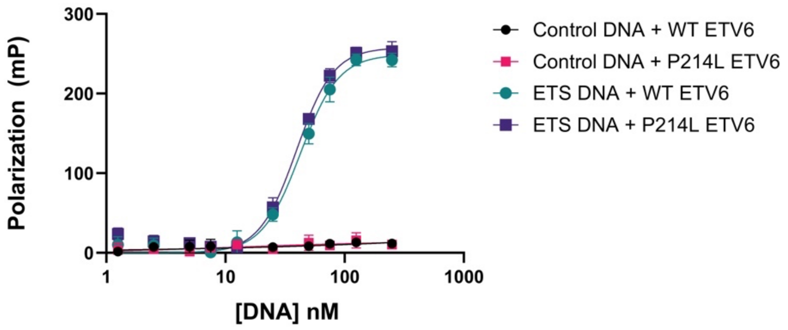


**Figure S1. Recombinant wild-type and P214L ETV6 constructs exhibit equal affinity for DNA harboring ETS binding sites.**

A fluorescence polarization interaction assay to assess binding affinity of full-length wild-type and P214L constructs to fluorescently-labeled DNA constructs. A DNA construct bearing an ETS binding site and a negative control are included. All ETV6 constructs harbor an A93D mutation to block PNT oligomerization, which is essential to isolate recombinant ETV6.

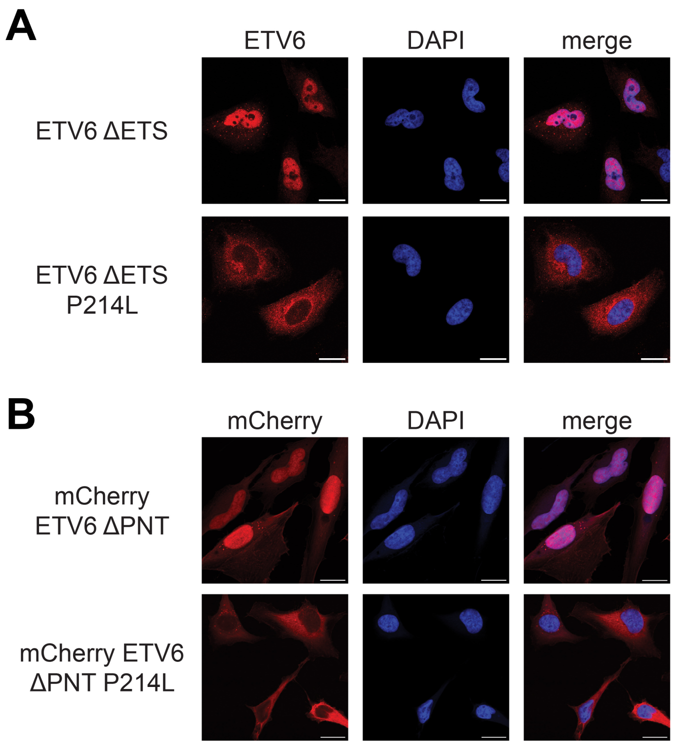


**Figure S2. P214L-mediated nuclear exclusion is maintained after removal of the ETS and PNT domains.**

(A) ETV6 immunofluorescence microscopy images of HeLa cells bearing stably integrated copies of the indicated ETV6 construct. Scale bar = 10 μm.

(B) Fluorescence confocal microscopy images of HeLa cells bearing stably integrated copies of the indicated mCherry-fused ETV6 construct. Scale bar = 10 μm.

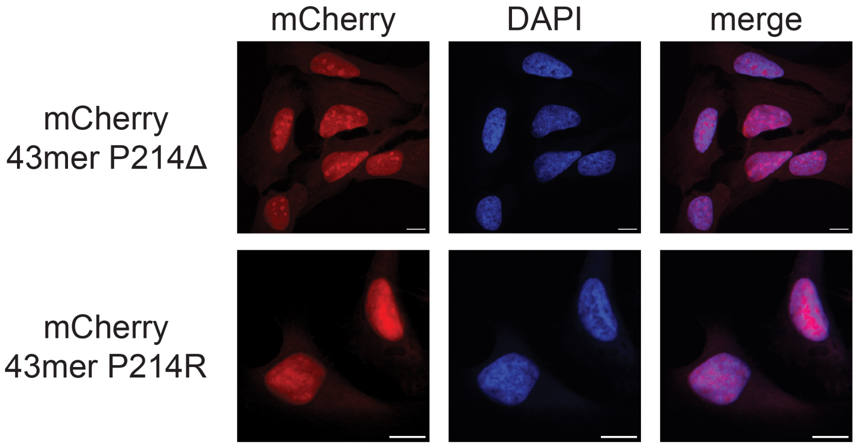


**Figure S3. The ETV6 P214 mutant mislocalization phenotype is specific to a leucine substitution.**

Fluorescence confocal microscopy z-stack images of HeLa cells bearing stably integrated copies of the indicated mCherry-fused ETV6 construct. Scale bar = 10 μm.

| Protein Name | Position | Sequence | Score |
|---|---|---|---|
| >ETV6 WT | 29-43 | SSTPLHVPVPRALRM | 0.060 |
| >ETV6 WT | 65-79 | DVAQWLKWAENEFSL | 0.017 |
| >ETV6 WT | 265-279 | VIQLMPSPIMHPLIL | 0.438 |
| >ETV6 WT | 306-320 | LSHREDLAYMNHIMV | 0.031 |
| >ETV6 WT | 308-322 | HREDLAYMNHIMVSV | 0.089 |

| Protein Name | Position | Sequence | Score |
|---|---|---|---|
| >ETV6 P214L | 29-43 | SSTPLHVPVPRALRM | 0.052 |
| >ETV6 P214L | 65-79 | DVAQWLKWAENEFSL | 0.017 |
| >ETV6 P214L | 200-214 | PLRSPLDNMIRRLSL | 0.611 |
| >ETV6 P214L | 265-279 | VIQLMPSPIMHPLIL | 0.240 |
| >ETV6 P214L | 306-320 | LSHREDLAYMNHIMV | 0.031 |
| >ETV6 P214L | 308-322 | HREDLAYMNHIMVSV | 0.089 |

**Figure S4. The NES prediction software LocNES identifies a strong NES from residues 200-214 in the ETV6 P214L construct.**

Full-length wild-type and P214L mutant ETV6 constructs were employed as input sequences in the LocNES prediction program. Scores > 0.2 are considered strong NES candidates. Note the NES from 265-279 contains multiple proline residue which are now known to be incompatible with XPO1 binding.

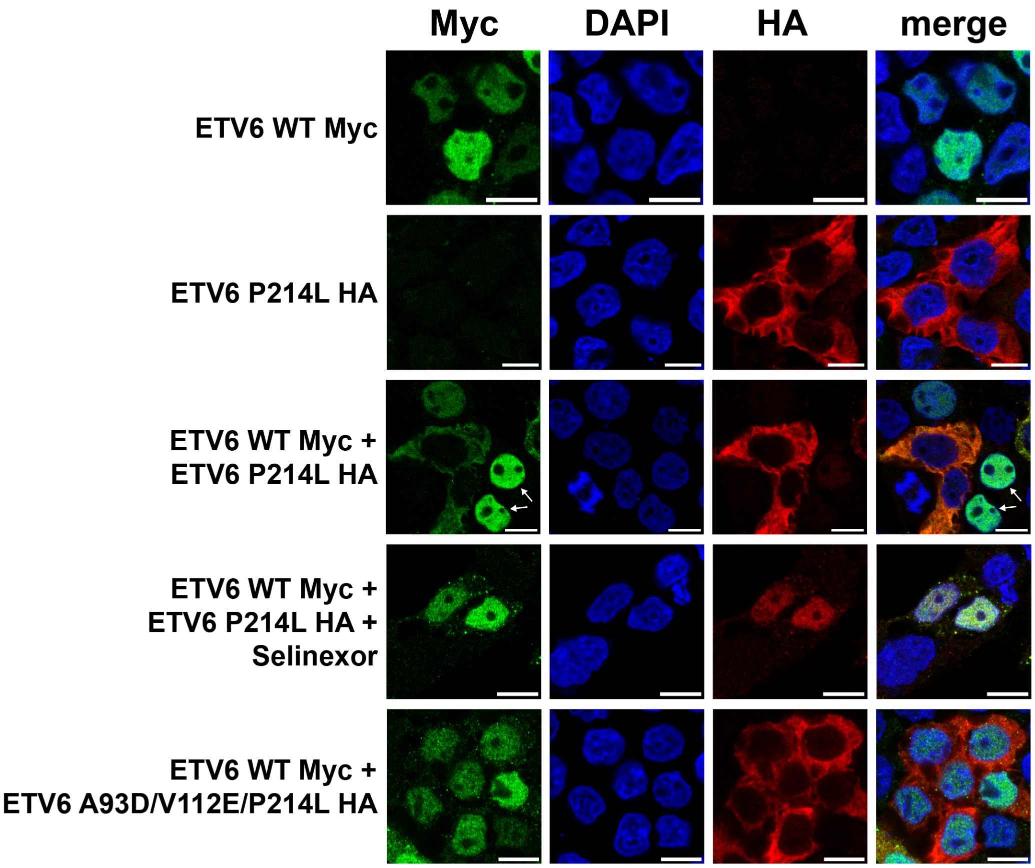


**Figure S5. PNT domain-mediated oligomerization leads to XPO1-dependent nuclear export of both mutant and wild-type ETV6 copies.**

Immunofluorescence microscopy images of HeLa cells bearing the indicated exogenous ETV6 construct(s). In the 'ETV6 WT Myc + ETV6 P214L HA' sample, the white arrows indicate cells containing only Myc-tagged wild-type ETV6. As expected, wild-type ETV6 is only mislocalized in cells expressing both wild-type and mutant constructs. Note: the A93D and V112E mutations block PNT domain oligomerization. Scale bar = 10 $\mu$m.

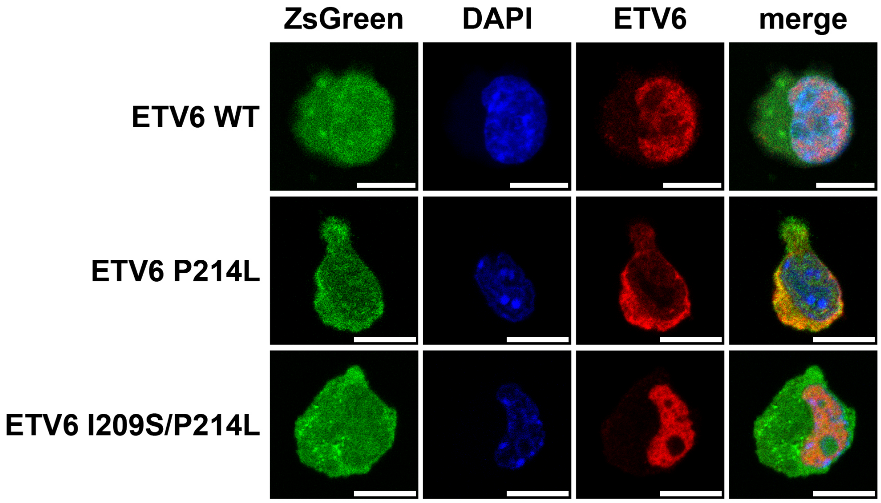


**Figure S6. ETV6 P214L-dependent mislocalization in Ba/F3$^{NRAS}$ cells.**

Immunofluorescence microscopy images of Ba/F3$^{NRAS}$ cells bearing the indicated stably incorporated ETV6 construct. Note: All ETV6 transcripts contain an IRES-ZsGreen element to indicate expression of the exogenous construct. Scale bar = 10 μm.

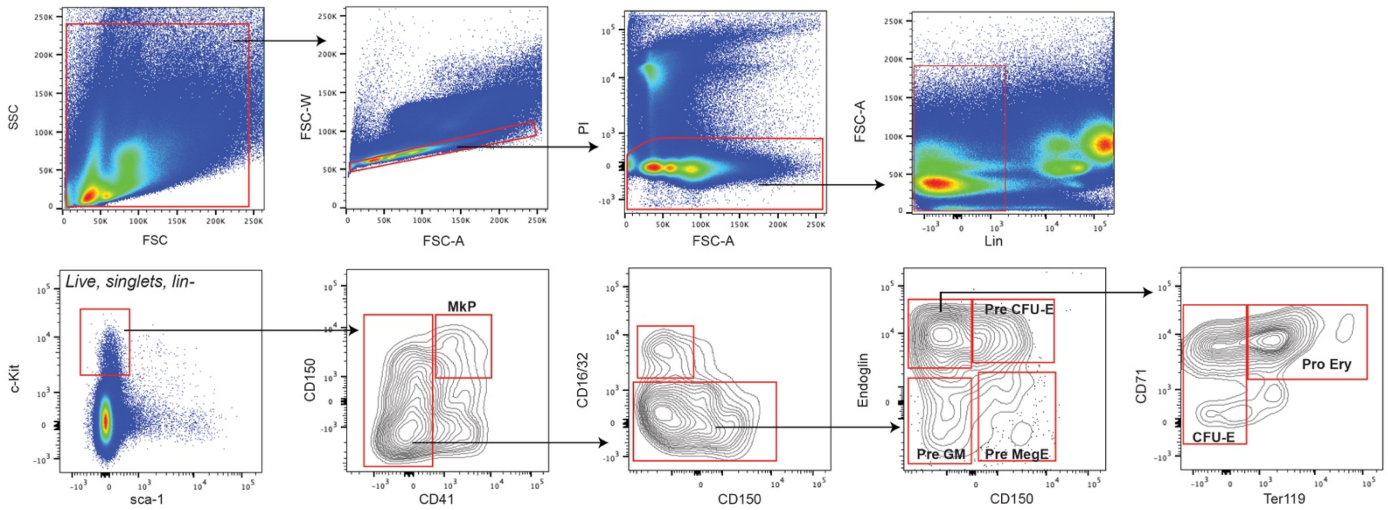


**Figure S7. The flow cytometry gating strategy for the myeloerythroid progenitor cell hierarchy.**

For full summary of gating protocol, see Material and Methods.

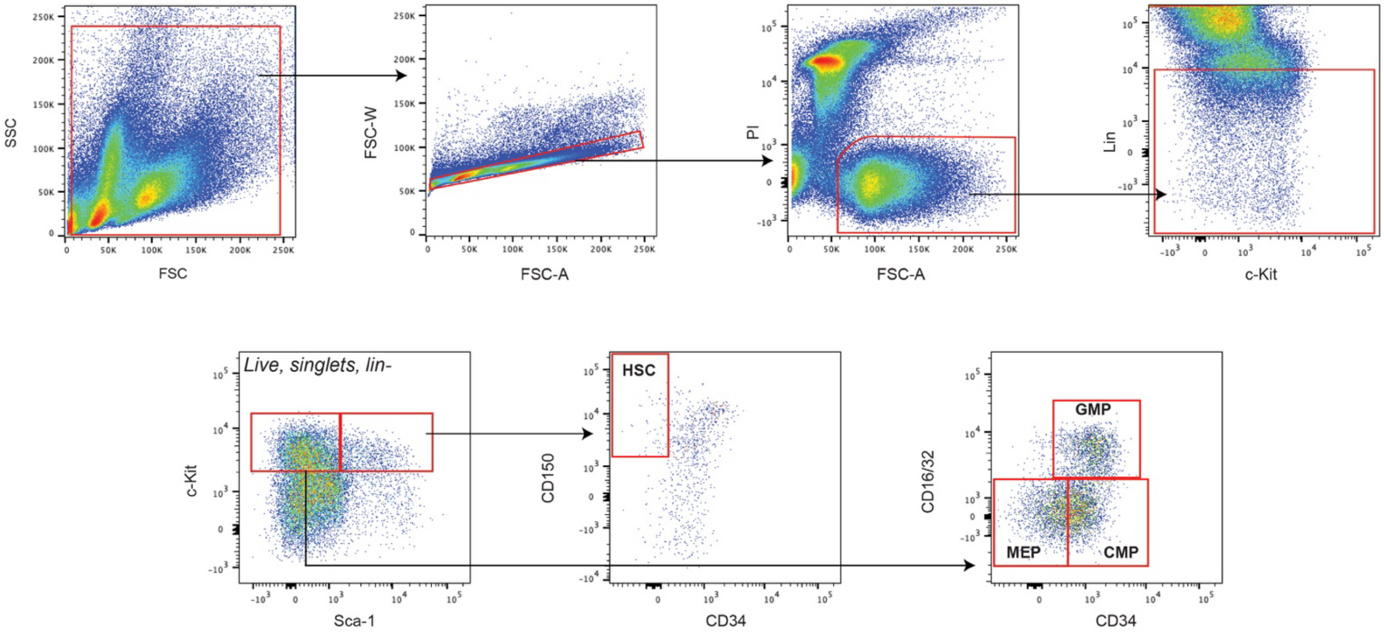


**Figure S8. The flow cytometry gating strategy for hematopoietic stem and progenitor cell populations.**

For full summary of gating protocol, see Material and Methods.

A

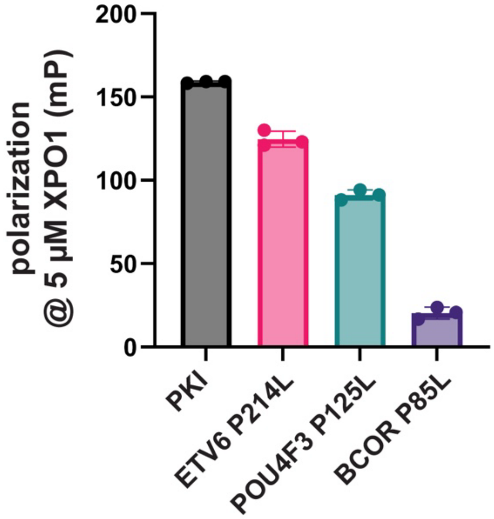


B

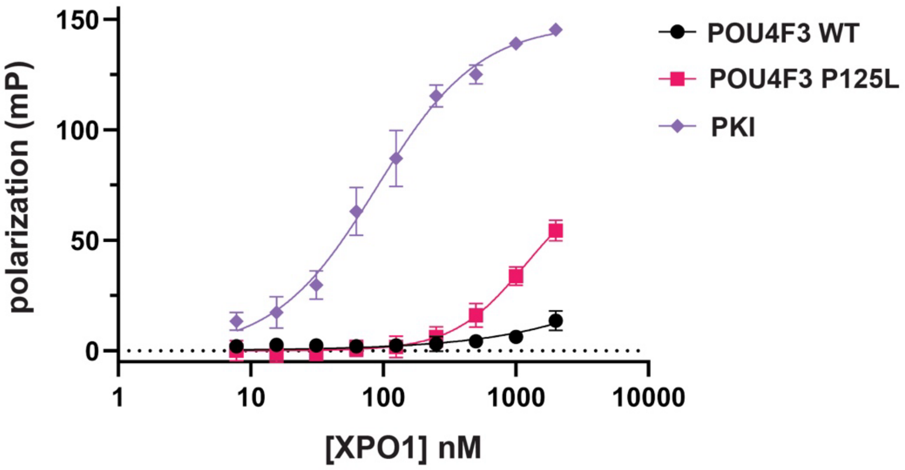


**Figure S9. Reconstituted XPO1:RanGTP interaction assay with various pathogenic variant 'hits' from the computational screen for mutation-dependent NESs.**

(A) A fluorescence polarization interaction assay to assess binding of synthetic, fluorescently-labeled mutant fragments to a reconstituted XPO1:RanGTP nuclear export receptor complex. A fluorescently labeled variant of the strong XPO1 ligand peptide PKI is included. All datapoints include 5 $\mu$M XPO1 and 5 nM peptide. Error bars represent S.D. from n = 3 experimental replicates.

(B) A fluorescence polarization interaction assay to assess binding of synthetic, fluorescently-labeled mutant and wild-type POU4F3 protein fragments (amino acids 114-134) to a reconstituted XPO1:RanGTP nuclear export receptor complex. A fluorescently labeled variant of the strong XPO1 ligand peptide PKI is included. All datapoints include 5 nM peptide. Error bars represent S.D. from n = 3 experimental replicates.

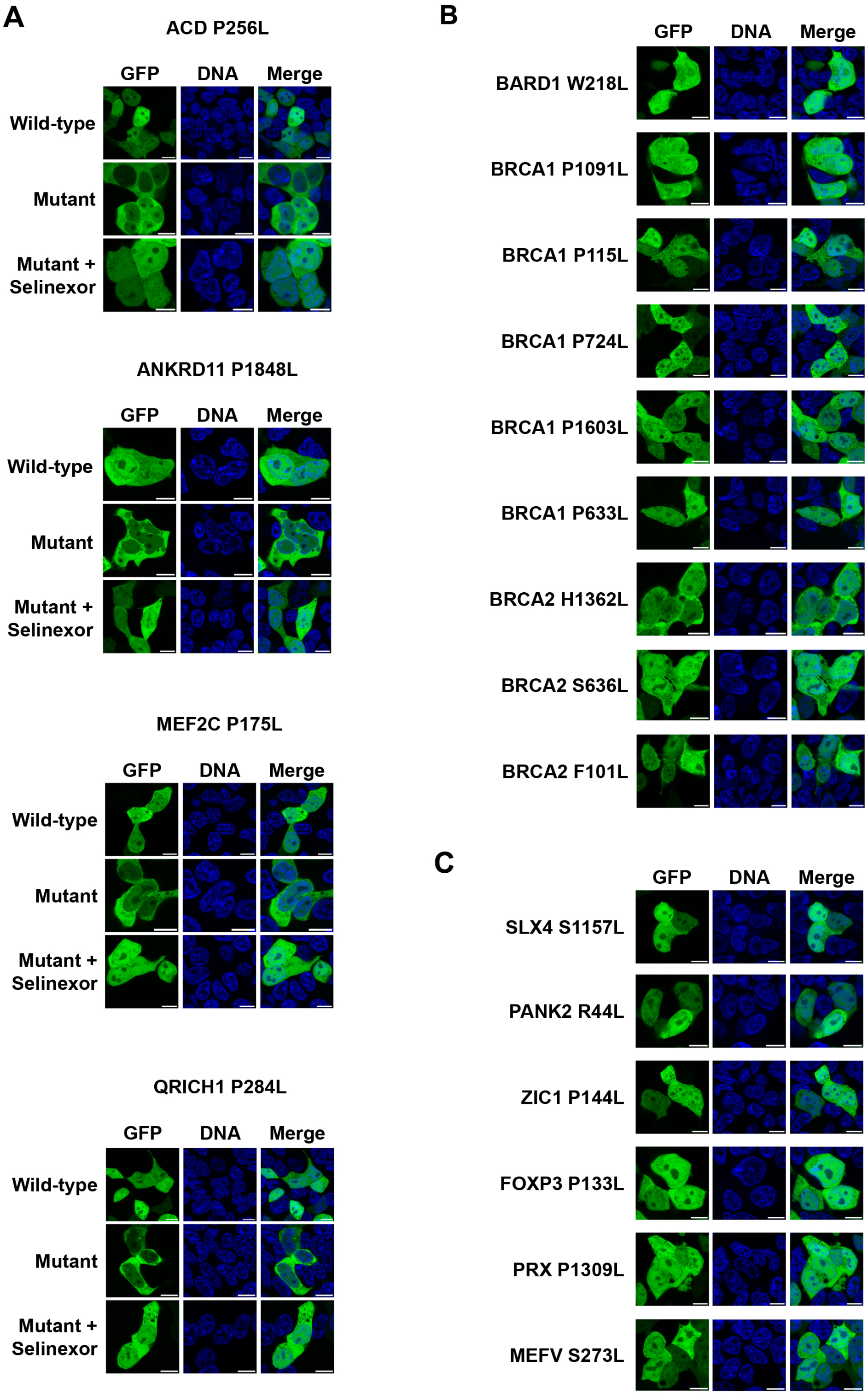


**Figure S10. A GFP-fused protein fragment approach to characterize mutation-dependent NES activity of computational screening 'hits'.**

(A) Fluorescence confocal microscopy images of live HEK293T cells bearing GFP-fused ACD (amino acids 243-273), ANKRD11 (amino acids 1828-1858), MEF2C (amino acids 157-187), and QRICH1 (amino acids 269-299) fragments from mutant proteins and wild-type counterparts. All mutations cause XPO1-dependent mislocalization. Images post-Selinexor treatment for 2 h are indicated. Scale bar = 10 μm.

(B) Fluorescence confocal microscopy images of live HEK293T cells bearing the indicated GFP-fused construct. All mutations are in HR-associated genes and all have no impact on protein localization. Scale bar = 10 μm.

(C) Fluorescence confocal microscopy images of live HEK293T cells bearing the indicated GFP-fused construct. All mutations are in non-HR-associated genes and all have no impact on protein localization. Scale bar = 10 μm.
